# STRIPAK associates with Rabex5-Rabaptin5 to promote Rab5-dependent early endosome formation and EGFR transport

**DOI:** 10.64898/2026.10.28.762573

**Authors:** Lydia Wunderley, Gabrielle Parkinson, Simon Nándor Sugár, Hideyuki Shimizu, Paul Fullwood, Eleanor Wendy Trotter, Chiara Francavilla, Philip Woodman

**Affiliations:** Faculty of Biology, Medicine and Health, University of Manchester, UK; Department of Biotechnology and Biomedicine, Technical University of Denmark; Cancer Research UK Manchester Institute, The University of Manchester, UK

**Keywords:** Strip1, SLMAP, Rab5-GEF, endocytosis, epidermal growth factor

## Abstract

His Domain Protein Tyrosine Phosphatase (HD-PTP) is required for multivesicular body (MVB) formation and endosome maturation, with Rab5-GTP levels raised upon HD-PTP depletion (Parkinson et al., 2021. doi: 10/1242/jcs.259192). Mass Spectrometry-based interactomics revealed that HD-PTP depletion increased association of Rabaptin5, a component of the early endosome Rabex5-Rabaptin5 Rab5-GEF, with multiple components of STRIPAK (striatin-interacting phosphatase and kinase). Native IPs and co-expression studies confirmed association between STRIPAK and Rabex5-Rabaptin5, including Strip1-Rabex5 and SLMAP-Rabaptin5 interactions. Consistent with functional Rabaptin5-STRIPAK links, co-depletion of Strip1 or Rabaptin5 with HD-PTP restored normal early endosome positioning and Rab5 distribution. Strip1 depletion impaired formation of central, Rab5-positive early endosomes, reduced cellular Rab5-GTP levels, and prevented the accumulation of internalised EGF-EGFR into centrally located endosomes. SLMAP depletion accentuated the central clustering of early endosomes and other endosomal compartments while also retarding the generation of new, EGFR-containing endosomes and EGFR lysosomal transport. We conclude that STRIPAK is a novel co-assembly factor of Rabex5-Rabaptin5, exerting subunit-specific effects on early endosomes.

## Introduction

Endocytosis monitors and modulates the plasma membrane proteome: internalised membrane proteins are delivered to early endosomes (EEs) and from there recycle to the cell surface or are sorted to towards lysosomal degradation (Cullen and Steinberg, 2018; Norris and Grant, 2020; Weeratunga et al., 2020). Recycling cargo is recognised by retrieval complexes that bind peptide-based sorting motifs and coordinate the formation of tubular recycling compartments (Carosi et al., 2023; Cullen and Steinberg, 2018; Norris and Grant, 2020; Weeratunga et al., 2020). The signal for lysosomal degradation is ubiquitination, which is recognised by Endosomal Sorting Complexes Required for Transport (ESCRT) complexes (Bishop et al., 2002; Christ et al., 2016; Henne et al., 2013; Piper et al., 2014; Raiborg et al., 2002; Rusilowicz-Jones et al., 2022). The ESCRT pathway sorts degradative cargo, exemplified by EGFR activated by saturating levels of EGF(Futter et al., 1996; White et al., 2006), into intralumenal vesicles within the developing multivesicular body (MVB) (Frankel and Audhya, 2018; Gatta and Carlton, 2019; Pfitzner et al., 2021; Quinney et al., 2019; Vietri et al., 2020).

EEs/MVBs are defined by the small GTPase Rab5 (Cezanne et al., 2020; Wandinger-Ness and Zerial, 2014; Zeigerer et al., 2012). Rab5-GTP recruits many important EE effectors. These include the membrane tethering factor early endosome antigen 1 (EEA1), which supports EE-EE homotypic fusion (Christoforidis et al., 1999a; Murray et al., 2016; Simonsen et al., 1998). Rab5-GTP also recruits the phosphatidylinositol-3 kinase (PI3 kinase), Vps34, which generates the EE-specific lipid phosphatidylinositol-3 phosphate (PI3P) (Christoforidis et al., 1999b). PI3P is recognised by EEA1 (Simonsen et al., 1998), members of the sorting nexin (SNX) family (Carlton et al., 2005; Stenmark and Aasland, 1999) that drive formation of recycling tubules (Carlton et al., 2004; Shortill et al., 2022; van Weering et al., 2012; van Weering et al., 2010), and the ESCRT-0 component Hrs (Raiborg et al., 2001), which initiates the MVB sorting pathway (Raiborg et al., 2002). Hence, Rab5 directly and/or indirectly promotes many aspects of EE behaviour.

Rab5-GTP is generated by a family of Rab5 guanine nucleotide exchange factors (Rab5-GEFs), each containing a catalytic Vps9 domain (Carney et al., 2006; Delprato and Lambright, 2007). The major Rab5-GEF acting at the EE/MVB is Rabex5/RabGEF1 (Horiuchi et al., 1997; Langemeyer et al., 2014), which forms a complex with Rabaptin5/RABEP1 (Lippe et al., 2001; Stenmark et al., 1995; Zhang et al., 2014). Rabex5 is activated allosterically by ubiquitin (Lauer et al., 2019; Mattera and Bonifacino, 2008; Mattera et al., 2006), ensuring that Rab5-GTP levels are maintained when ubiquitinated cargo is present in the EE (Ott et al., 2025). As MVB sorting progresses, cargo is deubiquitinated and enters ILVs, thereby reducing local ubiquitin levels (Ott et al., 2025; Rusilowicz-Jones et al., 2022).

Once endosomal sorting is complete, the MVB matures into a late endosome (LE). Endosomal maturation is governed by a process termed Rab conversion, whereby Rab5-GTP is replaced by Rab7-GTP (Huotari and Helenius, 2011; Podinovskaia and Spang, 2018; Rink et al., 2005). Rab conversion involves activation of Rab7 by the Rab7-GEF Mon1/Ccz1, which detects both Rabex5 and increasing levels of PI3P on the EE membrane. Alongside Rab conversion, PI3P is replaced by PI(3,5)P_2_. Rab7-GTP and PI(3,5)P_2_ recruit the HOPS complex, promoting late endosome-lysosome fusion (Borchers et al., 2021; Langemeyer et al., 2020; Nordmann et al., 2010; Poteryaev et al., 2010; Scott et al., 2014).

As well as Rab7 activation, Rab conversion entails reduced Rab5-GTP production (Rink et al., 2005). In part, this results from the fall in ubiquitin levels in EEs as MVB sorting is completed (Ott et al., 2025). However, the ESCRT scaffold His Domain Protein Tyrosine Phosphatase (HD-PTP) (Doyotte et al., 2008; Tabernero and Woodman, 2018) also provides a direct link between MVB sorting and Rab5-GTP levels. HD-PTP binds to multiple ESCRTs, including the ESCRT-0 component STAM2, the ESCRT-I components UBAP1 and TSG101, and the ESCRT-III component CHMP4B (Ali et al., 2013; Doyotte et al., 2008; Gahloth et al., 2017a; Gahloth et al., 2016; Gahloth et al., 2017b; Ichioka et al., 2007; Stefani et al., 2011). HD-PTP supports movement of MVB cargo along the ESCRT pathway (Doyotte et al., 2008; Kharitidi et al., 2015; Ma et al., 2015), by virtue of a shared binding site for STAM2 and CHMP4B within the HD-PTP Bro1 domain that promotes ESCRT exchange on HD-PTP (Ali et al., 2013). This site also binds directly to Rabaptin5 (Parkinson et al., 2021), suggesting mutually exclusive control of MVB sorting and Rab5-GTP production by HD-PTP. Indeed, alongside disruption of MVB sorting, depletion of HD-PTP prevents endosomal maturation and increases Rab5-GTP levels, effects that are reversed by co-depletion of Rabaptin5 (Parkinson et al., 2021). HD-PTP depletion also resulted in hyperphosphorylated Rabaptin5, highlighting a functional link between the proteins (Parkinson et al., 2021). The link between HD-PTP and Rabaptin5 appears central for endosomal maturation. To explore how HD-PTP controls Rab5-GTP production to influence Rab conversion, we examined how Rabaptin5 biochemistry is altered upon HD-PTP depletion. Here we find that HD-PTP depletion enhances interaction of the striatin-interacting protein phosphatase and kinase (STRIPAK) complex with Rabaptin5, uncovering STRIPAK (Hwang and Pallas, 2014; Kück and Pöggeler, 2024; Kuck et al., 2019) as a novel regulator of Rabex5-Rabaptin5 and EE function.

## Results

### HD-PTP depletion enhances association of multiple STRIPAK subunits with Rabaptin5

To understand how HD-PTP depletion enhances Rabaptin5-dependent Rab5 activation, we performed Mass Spectrometry (MS)-based interactomics using SILAC-labelled HeLa cells (see Fig. 1a for experiment and analysis flow chart), comparing native Rabaptin5 immunoprecipitates (IPs) from mock-depleted and HD-PTP-depleted HeLa cells (Table S1). Cell lysates were also subjected to proteomics analysis, to normalise Rabaptin5 IP data and to determine proteome-level effects of HD-PTP depletion (Table S2).

**Figure 1.**
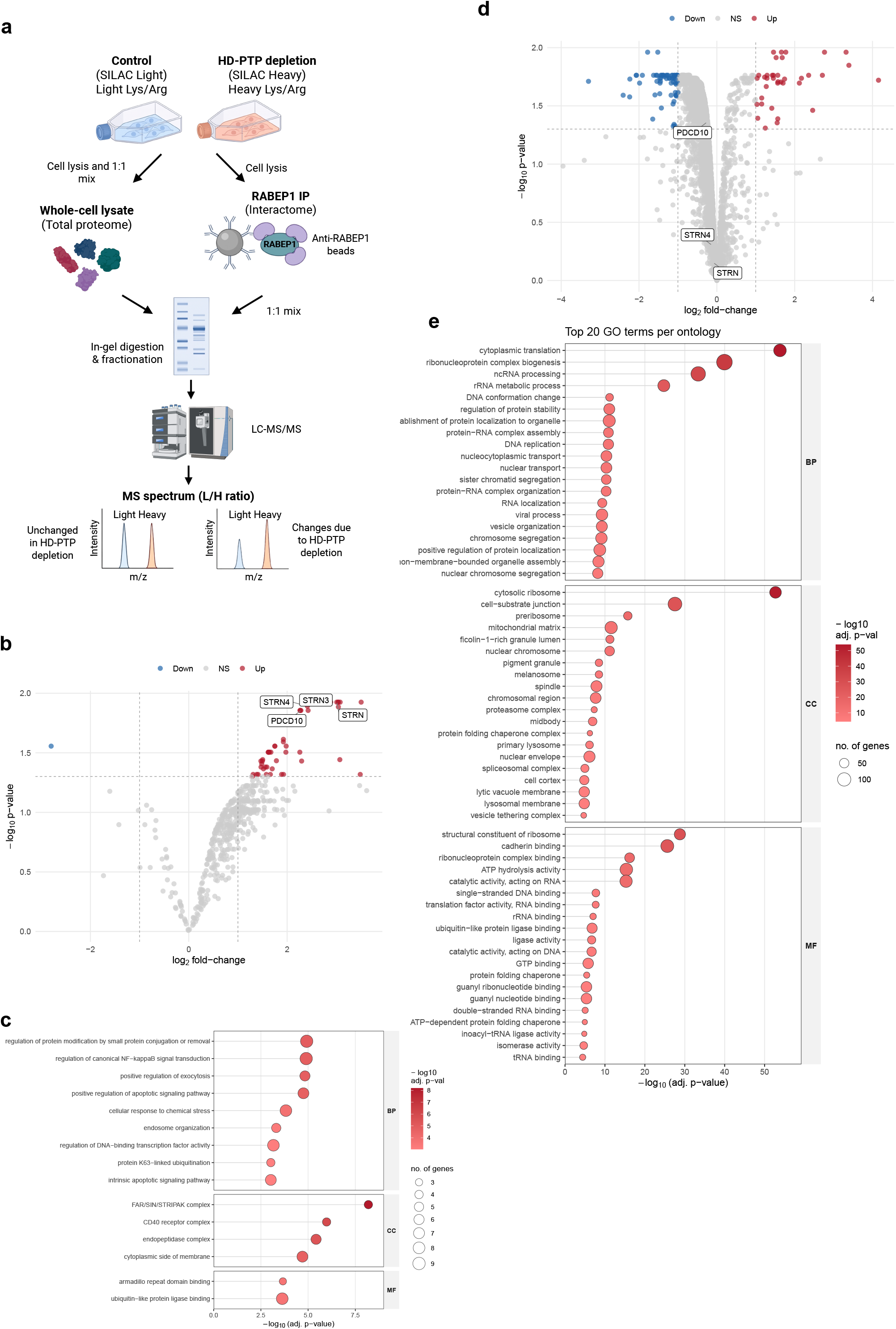
The STRIPAK complex associates with Rabaptin5 and binding is enhanced by HD-PTP depletion. **a:** Schematic showing SILAC-based MS analysis of Rabaptin5 interactors and proteome from control-vs HD-PTP-depleted cells. **b:** Volcano analysis of Rabaptin5 native IPs from control- and HD-PTP-depleted cells. Interactors with binding to Rabaptin5 significantly enhanced (red) or diminished (blue) upon HD-PTP depletion are highlighted. STRIPAK components and other important early endosome effectors are highlighted. **c:** Lollipop plots show GO terms enriched in Rabaptin5 IPs upon HD-PTP depletion. **d:** Volcano analysis of proteomes from control- and HD-PTP-depleted cells. **e:** Lollipop plots show GO terms enriched in proteome upon HD-PTP depletion.

Validation experiments confirmed that Rabaptin5 was immunoprecipitated by anti-Rabaptin5 specifically and with similar efficiency from control and HD-PTP-depleted cells (Fig. S1a). Density normalisation analysis (Fig. S1b) and pair-wise sample-sample correlation analysis (Fig. S1c) determined that IP1 and IP3 were very closely matched whereas IP2 had somewhat lower protein levels. However, downstream bait normalisation analysis successfully identified which proteins in IP2 followed the bait patterns of IP1 and IP3 (Fig. S1d), allowing high-confidence integration of the 3 IP datasets. QC of the proteome data (Fig. S1e-g) showed close correlation between the 3 replicates, based on the number of proteins identified between replicates, the density plot before quantile normalisation, and pairwise sample-sample correlation. In all, 3671 proteins were found in all 3 replicates (Fig. S1e), which is consistent with previous publications (Francavilla et al 2016).

Volcano plot analysis revealed 50 proteins whose co-IP with anti-Rabaptin5 increased significantly, and one protein whose co-IP decreased significantly, by depletion of HD-PTP (Fig. 1b). Lollipop plot analysis (Fig. 1c) revealed key GO terms enriched in Rabaptin5 IPs from HD-PTP depleted cells. GO pathways (BP) included regulation of NF-kappaB signalling, regulation of exocytosis, endosome organisation, regulation of apoptotic signalling, and regulation of K63-linked polyubiquitination. Enriched interaction GO terms (MF) included armadillo repeat binding and ubiquitin-like protein ligase-binding. Strikingly, GO analysis of protein complexes (CC) revealed particularly strong enrichment in components of the conserved human striatin-interacting protein phosphatase and kinase (STRIPAK) complex (termed factor arrest [FAR] complex in S. cerevisiae (Frost et al., 2012; Pracheil and Liu, 2013)and septation initiation network [SIN] inhibitory phosphatase [SIP] complex in S. pombe (Frost et al., 2012; Simanis, 2015; Singh et al., 2011)).

STRIPAK comprises a scaffold containing 4 copies of striatin (STRN), STRN-3/SG2NA and/or STRN-4/zinedin; the catalytic PP2A subunit PP2AB/PPP2CB and PP2A scaffold subunit A alpha (PP2AA/PPP2R1A); multiple adaptor and linker proteins, including Strip1/FAM40A, CCM3/PDCD10, MOB4, SIKE1, and FGFR1OP2/SIKE2; and the membrane-anchored phospho-substrate adaptor protein, SLMAP (Couzens et al., 2013; Duhart and Raftery, 2020; Hwang and Pallas, 2014; Jeong et al., 2021; Kück and Pöggeler, 2024; Kuck et al., 2019; Tanti et al., 2023; Wigle et al., 1997). Each of these components was present in Rabaptin5 IPs from control cells and enriched in Rabaptin5 IPs from HD-PTP-depleted cells (Fig. 1b; Table S1). STRIPAK associates with multiple protein kinases, principally within the Geminal centre kinase (GCK) family (Bae et al., 2020; Caputo et al., 2023; Kück and Pöggeler, 2024). One of these, MST4/STK26, was present in Rabaptin5 IPs and enriched in IPs from HD-PTP-depleted cells (Fig. 1b; Table S1).

Other components notably enriched in Rabaptin5 IPs from HD-PTP-depleted cells included Rab5 and EEA1, consistent with the increased Rab5-GTP levels seen upon HD-PTP depletion (Parkinson et al., 2021) (Table S1). The ESCRT-0 components Hrs and STAM were also enriched (Table S1), consistent with ESCRT-0 being trapped within EE membrane domains upon HD-PTP depletion (Ali et al., 2013) and the recruitment of ESCRT-0 to PI3P-enriched membrane domains (Raiborg et al., 2001) generated downstream of Rab5-GTP (Cezanne et al., 2020; Zerial and McBride, 2001).

Differential proteome analysis by Volcano plot (Fig. 1d) followed by GO term (Fig. 1e) analysis revealed multiple pathway components were altered by HD-PTP depletion, with RNA and translation pathways surprisingly enriched, consistent with the enrichment of ribosome components also observed. Aside from these pathways, the most notable enrichment was for cell-substrate junction and cadherin complex components. Importantly, none of the proteins significantly enriched in Rabaptin5 IPs from HD-PTP-depleted cells was differentially present in the proteomes from control and HD-PTP-depleted cells (Fig 1d). Additionally, none of these 50 proteins (Fig. S1h) including all STRIPAK components (Fig. S1i) displayed any systematic enrichment in the proteome of HD-PTP-depleted cells between experimental replicates. Hence, altered expression could not explain the observed differences in co-IP efficiency of these components between experimental conditions. Rabex5 was present in Rabaptin5 IPs from control cells, and levels did not increase significantly upon HD-PTP depletion (Table S1). Hence, the increase in Rab5-GTP levels in HD-PTP-depleted cells is not simply a consequence of increased Rabex5-Rabaptin5 association.

Enhanced association of STRIPAK components with Rabaptin5 upon HD-PTP depletion was confirmed by Western blotting Rabaptin5 IPs for endogenous STRIPAK components. These showed that endogenous STRN-1, Strip1, SLMAP and PP2AA were all present in Rabaptin5 IPs from control cell lysates, and that their levels each increased significantly in Rabaptin5 IPs from HD-PTP-depleted cells (Fig. 2a-d). To further confirm association of multiple STRIPAK components with Rabaptin5, GFP-Rabaptin5 was co-expressed in HEK293 cells with a range of FLAG-tagged STRIPAK components. GFP-Rabaptin5 was pulled-down efficiently on anti-FLAG agarose beads when co-expressed with FLAG-SLMAP (Fig. 3a), suggesting this may be the primary point of interaction between Rabaptin5 and STRIPAK. HA-Rabex5 was pulled-down efficiently on anti-FLAG agarose beads when co-expressed with FLAG-Strip1, and FLAG-SIKE2 (Fig.3b). Altogether (Fig. 3c), these findings indicate that the association between Rabex5-Rabaptin5 and STRIPAK is complex, potentially involving multiple interactions.

**Figure 2.**
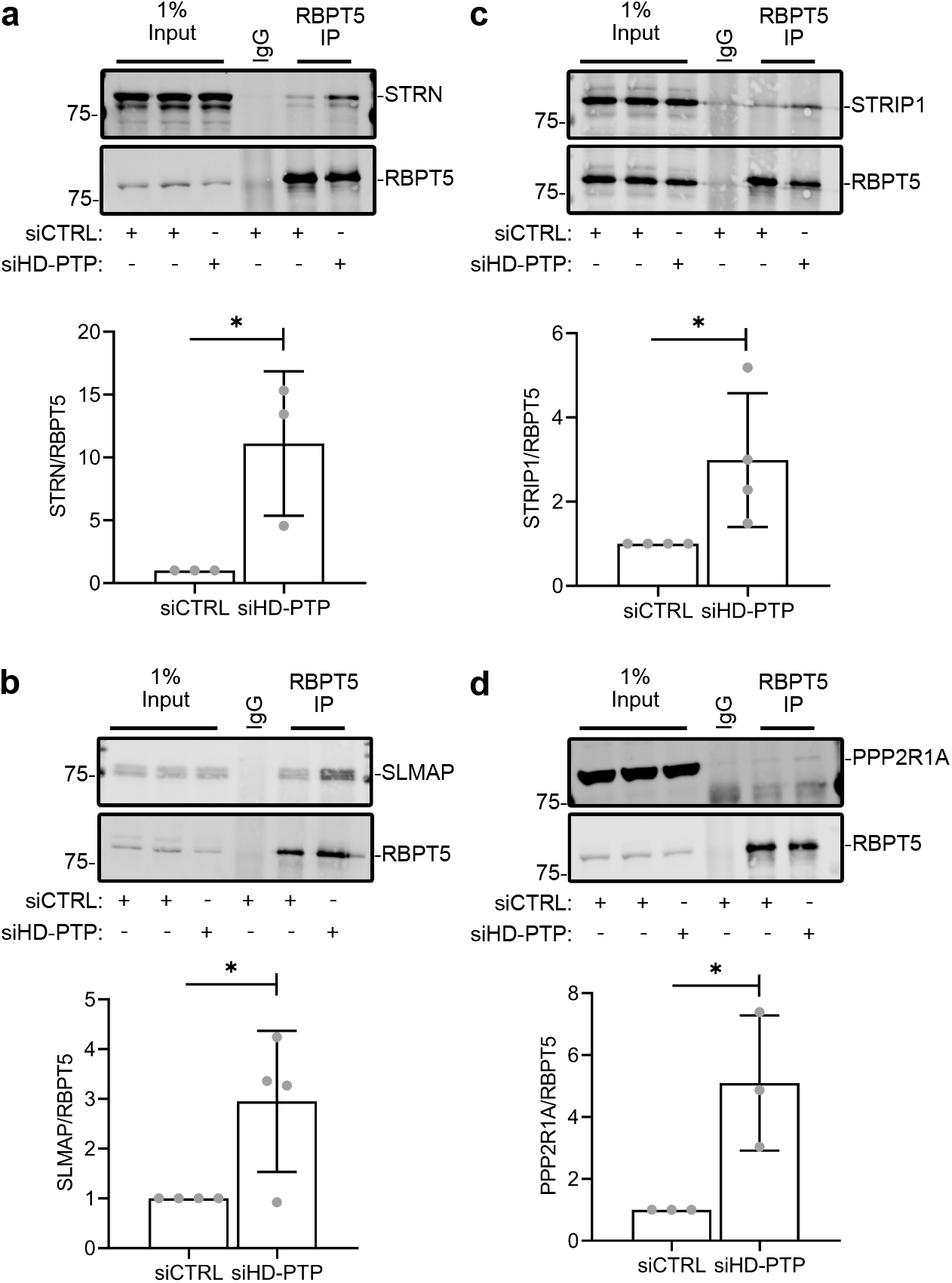
Native IPs confirm STRIPAK binding to Rabaptin5 is enhanced by HD-PTP depletion. **a-d:** Native Rabaptin5 IPs from control and HD-PTP-depleted HeLa cells Western blotted for striatin (**a**), SLMAP (**b**), Strip1 (**c**), PPP2R1A (**d**). Upper panels: representative blots. Lower panels quantification from independent experiments.

**Figure 3.**
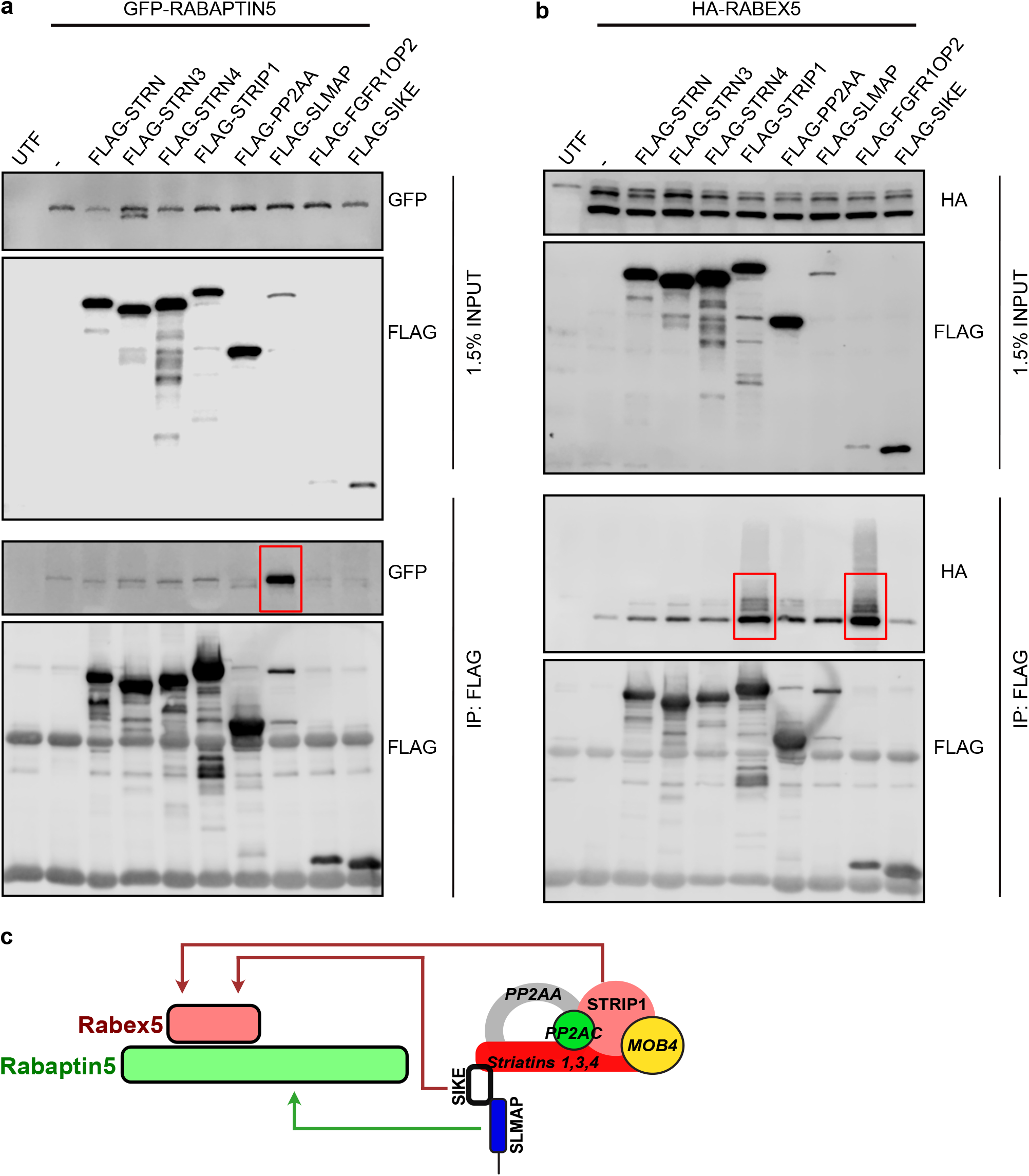
Multiple interactions between STRIPAK and Rabex5-Rabaptin5. Lysates from HEK293 cells co-expressing GFP-Rabaptin5 **(a)** or HA-Rabex5 **(b**) with a range of FLAG-tagged STRIPAK subunits were subjected to IP on FLAG-beads. Key interactions are highlighted with red boxes. **c**: Schematic summarising outcome of binding experiments.

MS-based analysis showed the Rabaptin5-related protein Rabaptin5β/RABEP2 was also present in Rabaptin5 IPs (Table S1). Like Rabaptin5 (Rabaptin5α), Rabaptin5β associates with Rabex5 and supports Rabex5-dependent Rab5-GEF activity (Gournier et al., 1998). However, Rabaptin5β is unlikely to contribute to the increased co-IP of STRIPAK with anti-Rabaptin5 upon HD-PTP depletion. First, the Bro1-coiled-coil fragment of HD-PTP (HA-Bro1CC) interacts with GFP-Rabaptin5α but not with GFP-Rabaptin5β, as revealed by co-IPs from co-transfected cells (Fig. S2a). Second, GFP-Rabaptin5α strongly, but GFP-Rabaptin5β only weakly, co-localised with HA-Bro1CC when co-expressed (Fig. S2b). Third, while GFP-Rabaptin5α strongly interacted with FLAG-SLMAP, no comparable interaction was observed between GFP-Rabaptin5β and SLMAP (Fig. S2c). Hence Rabaptin5β binds neither HD-PTP nor SLMAP.

### Co-depletion of Strip1 rescues Rab5-endosome behaviour in HD-PTP-depleted cells

Depletion of HD-PTP disrupts MVB sorting of EGFR and other cargo (Doyotte et al., 2008; Kharitidi et al., 2015; Ma et al., 2015; Parkinson et al., 2015), prevents lysosome-bound cargo exiting Rab5-rich EEs (Parkinson et al., 2021), and generates perinuclear enlarged and often clustered EEs with high Rab5 content (Parkinson et al., 2021). Since these phenotypes correlate with increased association between Rabaptin5 and STRIPAK, we tested whether co-depletion of STRIPAK components might restore normal EE positioning in HD-PTP-depleted cells.

RNAi-based depletion of Strip1 in HeLa cells achieved ∼50-60% reduction in Strip1 levels (Fig. S3a). We could not deplete Strip1 levels further, presumably because Strip1 is an essential protein. However, co-depletion of Strip1 rescued the EE enlargement/clustering phenotype observed upon HD-PTP depletion, similarly to co-depletion of Rabaptin5 (Fig. S3b for assessment of Rabaptin5 KD), as evidenced by staining cells with the EE marker EEA1 (Fig. 4a,b). In contrast, co-depletion of SLMAP (Fig. S3a for assessment of SLMAP KD) did not (Fig. 4a,b). Unfortunately, we could not assess the impact of combining depletion of STRN-1, -3 and -4 with HD-PTP depletion, since we found that combining STRN-1, -3 and -4 siRNAs was cytotoxic in HeLa cells.

**Figure 4.**
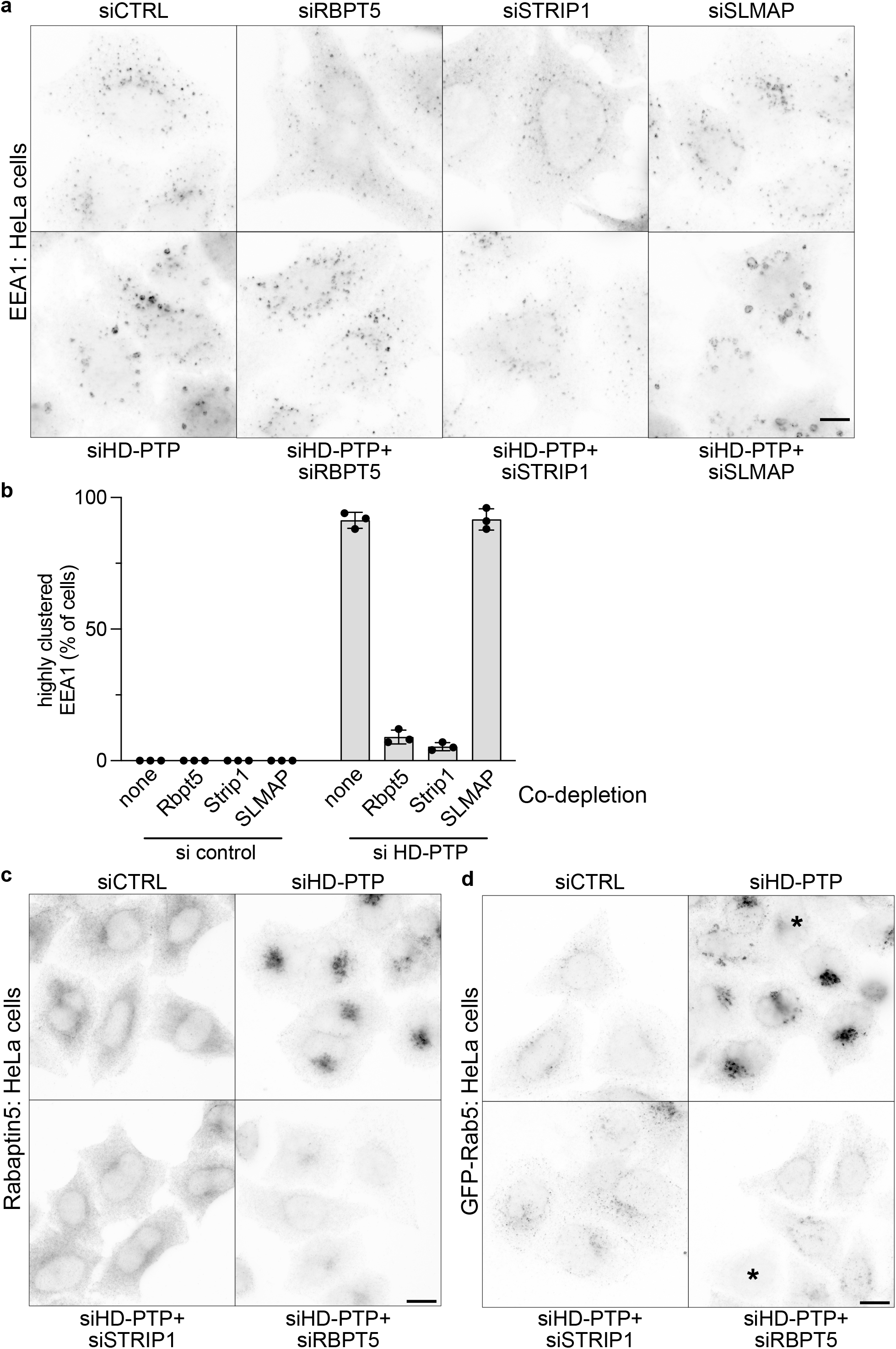
Strip1 or Rabaptin5 co-depletion rescues impact of HD-PTP depletion on early endosomes. HeLa cells were depleted using siRNAs as indicated and stained for EEA1. **a:** representative images, **b:** quantitation of phenotypes from 3 independent experiments. **c:** HeLa cells were depleted using siRNAs as indicated and stained for Rabaptin5. **d:** HeLa cells were depleted using siRNAs as indicated, transduced with GFP-Rab5 and stained for GFP. Scale bars = 10 µm.

Since Strip1 or Rabaptin5 co-depletion restored normal EE characteristics to HD-PTP-depleted cells, we assessed the impact of their co-depletion on the distribution of endogenous Rabaptin5. In control cells, Rabaptin5 was largely cytosolic, with somewhat increased labelling intensity in the perinuclear region evident (Fig. 4c). Upon HD-PTP depletion, Rabaptin5 localised strikingly to clustered, perinuclear membranes and the cytoplasmic pool of Rabaptin5 was diminished (Fig. 4c). These effects were reversed by co-depleting Strip1 (Fig. 4c). As expected, Rabaptin5 staining was essentially absent when Rabaptin5 was co-depleted (Fig. 4c). These effects were mirrored by the impact of Strip1 or Rabaptin5 co-depletion on Rab5 distribution, assessed by transducing cells with low levels of lentivirus-encoded GFP-Rab5 to best visualise all Rab5 pools: Rabaptin5 or Strip1 co-depletion eliminated the strong perinuclear concentration of GFP-Rab5 seen upon HD-PTP depletion (Fig. 4d). In summary, Strip1 co-depletion closely followed Rabaptin5 co-depletion in restoring normal EE and Rab5 positioning to HD-PTP-depleted cells. Unfortunately, we could not reliably assess the impact of Strip1 co-depletion on EGFR trafficking in HD-PTP-depleted cells, since Strip1 depletion slightly reduces levels of EGFR (see Fig. S3a) and these effects were compounded upon Strip1/HD-PTP co-depletion (data not shown).

### Strip1 is required to generate central Rab5-positive EEs containing internalised EGFR

These data suggest that Strip1 may be important for Rabex5-Rabaptin5 function in maintaining EEs. To test this, we depleted Strip1 and compared effects on EEs with Rabaptin5 depletion. In Rabaptin5- or Strip1-depleted HeLa cells, EEs had reduced levels of EEA1 compared to in control-depleted cells (Fig.5a,b). Centrally located EEs clearly labelled with antibody against endogenous Rab5 in control-depleted cells but labelling of EEA1-positive endosomes for Rab5 was less distinct in Strip1- or Rabaptin5-depleted cells (Fig. 5a). Quantitative analysis showed that EEs contained less EEA1 and Rab5 when Rabaptin5 or Strip1 were depleted (Fig. 5b). A reduction in endogenous Rab5 labelling on EEs was also observed in RPE-1 cells, which often contain a central, juxtanuclear arrangement of large EEA1-positive endosomes that stain strongly for Rab5 (Fig. S3c). Strip1 or Rabaptin5 depletion also caused EEs to become more dispersed relative to those in control-treated cells (Fig. 5a, Fig. S3c). The impact of Strip1 depletion on Rab5-positive endosomes in RPE cells was confirmed using lentivirus-encoded GFP-Rab5. In control cells, GFP-Rab5 localised to a central pool of EEA1-positive endosomes and to smaller endosomes located towards the cell periphery (Fig. S3d). In Strip1-depleted cells, the central concentration of EEA1-positive endosomes was diminished and GFP-Rab5 became more evenly distrusted throughout the cytoplasm (Fig. S3d).

**Figure 5.**
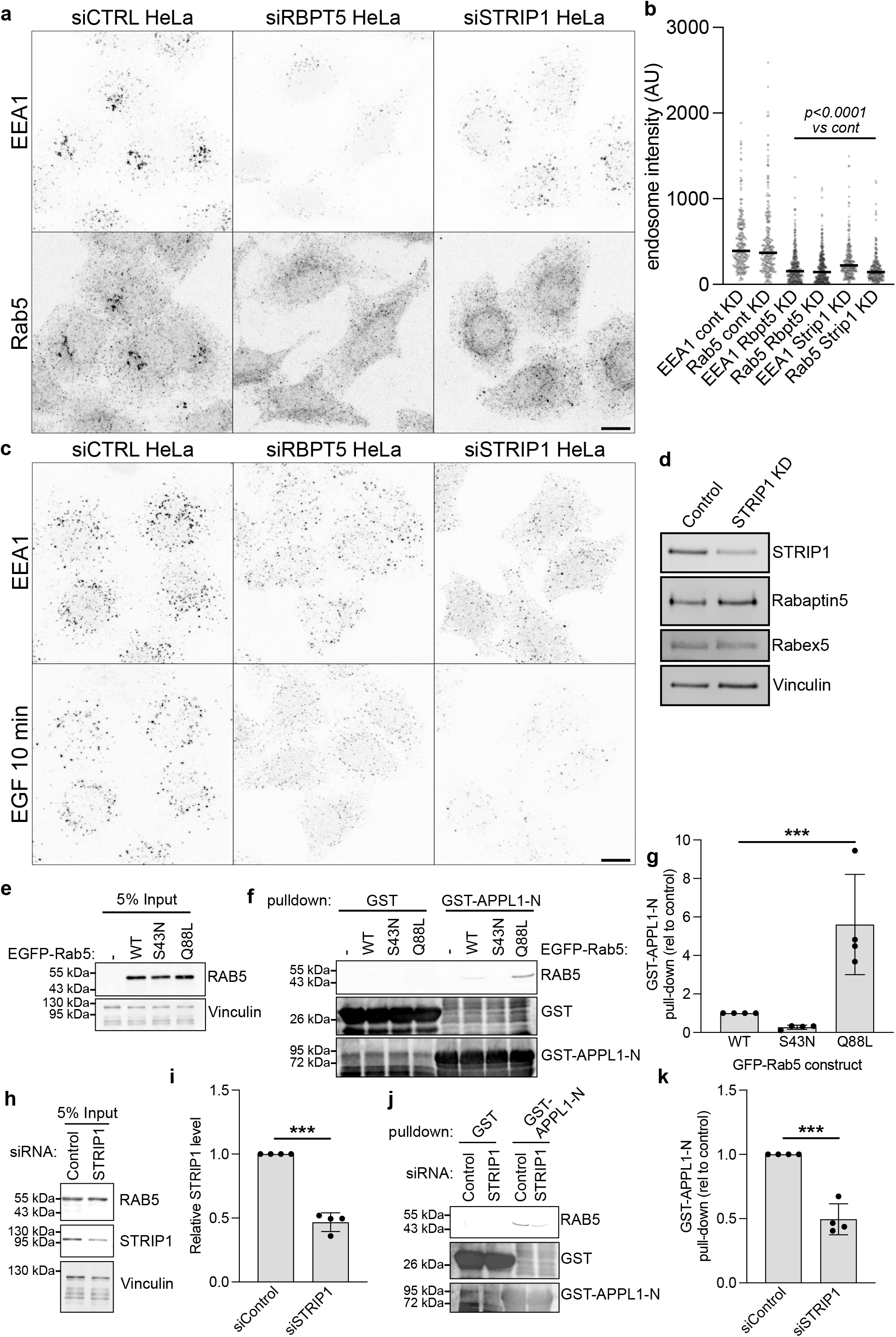
Strip1 is important for generating Rab5-early endosomes. **a:** HeLa cells were depleted using siRNAs as indicated and stained for EEA1. representative images. **b:** quantitation of EEA1 and Rab5 staining intensity from randomly selected endosomes. **c:** HeLa cells were depleted using siRNAs as indicated, bound and chased for 10min with ^A88^EGF and stained for EEA1. **d:** Western blot of Strip1 siRNA showing levels of Rabex5 and Rabaptin5. **e-k:** GST pull-down analysis of Rab5A activity using the N-terminal Rab5-binding domain of APPL1 (GST-APPL1-N), which preferentially associates with active, GTP-bound Rab5. Validation of the GST pull-down assay using Rab5A activity-state controls, showing representative input (**e**) and IP (**f**) Western blots, and quantitation from 4 independent experiments (**g**). Effect of STRIP1 depletion on Rab5A activity showing representative input Western blots (**h**), quantitation of blots from 4 independent experiments (**i**), representative IP Western blots (**j**) and quantitation of blots from 4 independent experiments. Scale bars = 10 µm.

We then tested the impact of Strip1 vs Rabaptin5 depletion on the accumulation of internalised EGF in EEA1-positive endosomes. Alexa-conjugated EGF was bound to cells for 1h at 4°C and internalised by warming cells to 37°C. In control-depleted HeLa cells, EGF rapidly entered EEA1-positive endosomes, including those in the juxtanuclear area (Fig. 5c). EGF-containing endosomes were less intensely labelled with EGF, contained less EEA1 and Rab5, and were more dispersed in Rabaptin5-depleted cells (Fig. 5c). This was also the case in Strip1-depleted cells (Fig. 5c), though the dispersal of EEs was more exaggerated, and the number of EGF-containing endosomes was diminished. This may be partly explained by the slightly reduced levels of EGFR in Strip1-depleted cells (Fig. S3a). In summary, Strip1 depletion closely phenocopies the impact of Rabaptin5 depletion of the behaviour of EEs. This is not a consequence of Strip1 depletion affecting Rabaptin5 or Rabex5 protein levels, since these were unaffected by Strip1 depletion (Fig. 5d; Fig. S3a). The impact of Strip1 depletion of inward movement of EGFR-containing endosomes confirms previous findings (Sakuma et al., 2014).

Altogether, the above findings suggest that Strip1 may be important for generating Rab5- and EEA1-positive endosomes, via effects on Rabex5-Rabaptin5. To test the impact of Strip1 depletion on Rab5-GTP levels, we employed a biochemical reporter. Here, lysates were made from cells expressing GFP-Rab5 and incubated with glutathione-S-transferase (GST)-tagged APPL1 N-terminal domain (GST-APPL1-N), which binds to activated Rab5 (Hiragi et al., 2022). Rab5-GTP levels were assessed by how much GFP-Rab5 was pulled down on glutathione beads. Control experiments showed that GST-APPL1-N pulled down WT GFP-Rab5 but not a dominant-negative GFP-Rab5 mutant, GFP-Rab5^S34N^, while pulling down more effectively a GTPase-defective Rab5 mutant, GFP-Rab5^Q88L^ (Fig. 5e-g). Strip1 depletion reduced pull-down of WT GFP-Rab5 by corresponding levels (Fig. 5h-k). Hence, Strip1 is important for producing/maintaining Rab5-GTP.

We then tested whether CRISPR-based depletion of Strip1 also affects endosomes. Using RPE-1 cells targeted with OsTIR1(F47G) to the AAVS safe harbour site, we used CRISPR-Cas9 gene editing to replace endogenous Strip1 with Strip1 containing a C-terminal sequence encoding an auxin-inducible degron (AID) and FK506-binding protein 12 carrying a F36V mutation (FKBP12^F36V^) (dTAG) (Belbelazi et al., 2026; Nabet et al., 2018). These sequences provide a double degron (DD) with the potential to rapidly degrade Strip1 by addition of either the auxin analogue 5-phenyl-indole-3-acetic acid (5-Ph-IAA) or dTAGV-1 (Nabet et al., 2018; Yesbolatova et al., 2020). Clonal selection of cells followed by sequencing of genomic DNA allowed us to generate 3 cell lines incorporating gene-edited Strip1-DD. Incorporation of Strip1-DD was confirmed by Western blotting for Strip1 (Fig. 6a). Levels of Strip1-DD in each line were slightly lower than levels of Strip1 in parental RPE-1 cells, suggesting a low level of degradation of Strip1-DD occurs even in the absence of induction. In each clonal line, Strip1-DD was completely degraded within 2h of addition of dTAGV-1 (Fig. 6a).

**Figure 6.**
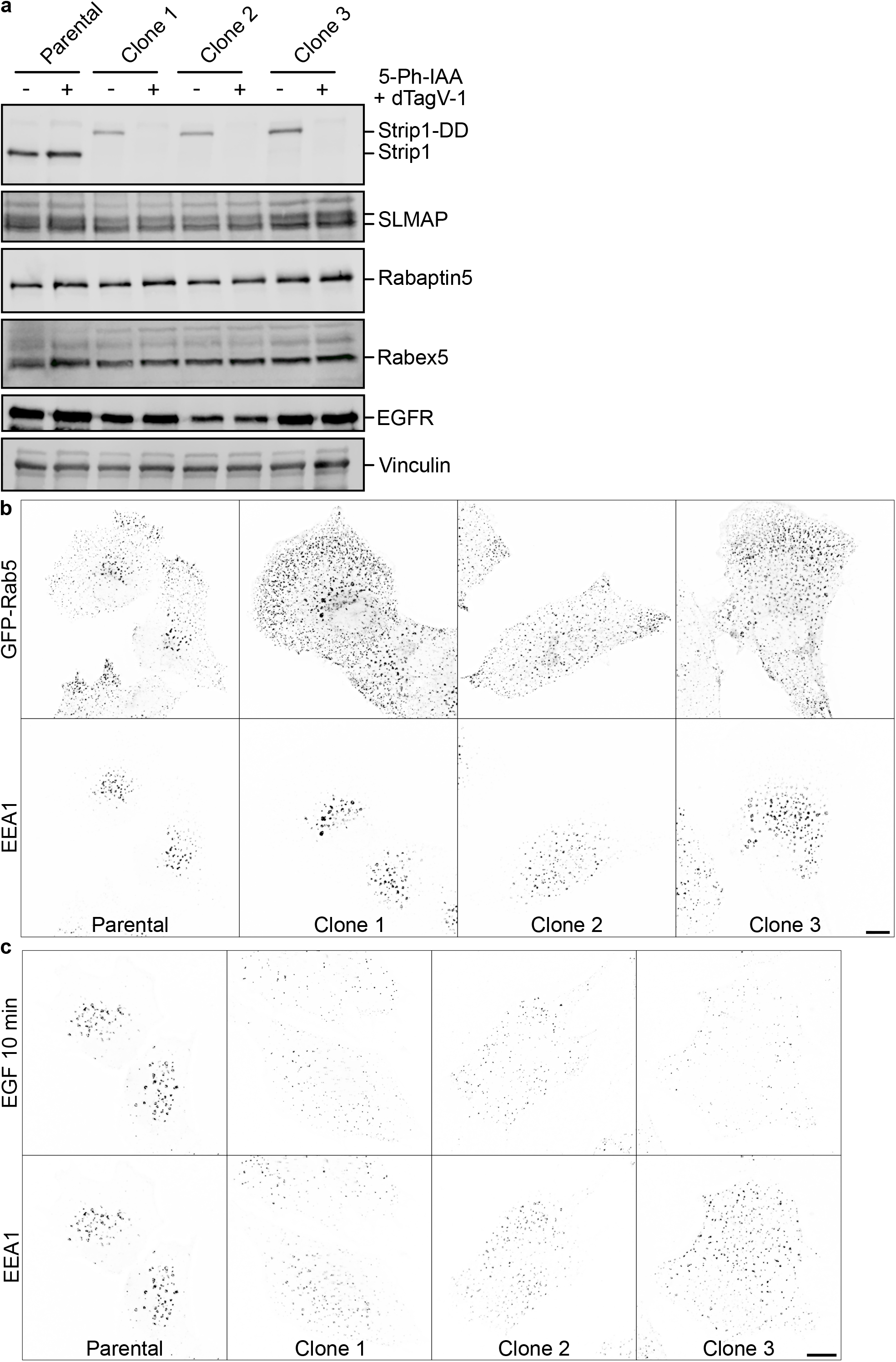
Strip1 degron cells display impaired early endosome formation. **a:** Western blot showing levels of Strip1 in 3 independent Strip1-DD clones vs parental RPE-1 cells, treated for 2 h with or without dTAGV-1. **b:** Parental RPE-1 cells or Strip1-DD cells were transduced with GFP-Rab5 and treated with dTAGV-1 for 4 h prior to staining for GFP and EEA1. **c:** Parental RPE-1 cells or Strip1-DD cells were bound and chased for 10min with ^A88^EGF and stained for EEA1. Scale bars = 10 µm.

Using GFP-Rab5 to visualise all pools of Rab5 showed that Rab5 distribution was strongly affected Strip1-DD clones compared to parental RPE-1 cells treated with dTAG-V1 (Fig. 6b). In parental RPE-1 cells, GFP-Rab5 concentrated in large juxta-nuclear EEA1-positive endosomes, as for endogenous Rab5 (Fig. S4c), but was also detected in a pool smaller, EEA1-negative endosomes at the cell periphery (Fig. 6b). In dTAGV-1-treated Strip1-DD cells, this polarised distribution of GFP-Rab5 was disrupted. While a centrally distributed pool of GFP-Rab5-positive, EEA1-positive endosomes was still evident (albeit somewhat less clustered than in parental cells), the pool of GFP-Rab5-positive, EEA1-negative endosomes was strongly enhanced in the periphery and more fully occupied the intermediate region. (Fig. 6b).

We then followed the uptake of Alexa^488^EGF bound to cells at 4°C followed by a chase at 37°C for 10 min. In parental RPE-1 cells treated with dTAG-V1, Alexa^488^EGF reached EEA1-positive endosomes, many centrally located, within 10 min. In contrast, in Strip1-DD cells acutely depleted of Strip1 by dTAG-V1, Alexa^488^EGF was internalised into small endosomes distributed throughout the cell periphery. Furthermore, the distribution of EEA1 labelling was strongly affected, with most EEA1 now occupying small EEs throughout the cell (Fig. 6c). In summary, acute depletion of Strip1 alters the steady-state distribution of Rab5-containing endosomes, impairs the inward redistribution of internalised EGF into new endosomes and reduces the overall amount of EGF reaching larger, EEA1-positive endosomes.

### SLMAP is important for organising endolysosomal compartments and forming EGFR-containing EEs

While Strip1 co-depletion rescues the impact of HD-PTP depletion on endosomes, SLMAP co-depletion does not (Fig. 4b,c), despite SLMAP associating robustly with Rabaptin5 (Fig. 3a). We therefore examined how SLMAP depletion affects endosomes. In contrast to RNAi-mediated Strip1 depletion, RNAi-mediated SLMAP depletion did not reduce the intensity of EEA1 or Rab5 labelling on endosomes in HeLa cells. Instead, the extent of EE clustering in the perinuclear region increased, and Rab5 staining became more concentrated in the perinuclear region (Fig. 7a). In contrast to Strip1 depletion, SLMAP depletion did not reduce levels of Rab5 on EEs in RPE-1 cells (Fig. 7b) and tended to increase EEA1 clustering. Furthermore, SLMAP depletion had contrasting effects to Strip1 depletion on GFP-Rab5 distribution. In SLMAP-depleted RPE-1 cells, centrally located EEs contained more EEA1 and GFP-Rab5 and were larger and more clustered than in control-depleted cells (Fig. 7c). Additionally, the population of peripheral GFP-Rab5-positive, EEA1-negative EEs was substantially diminished in SLMAP-depleted RPE-1 cells (Fig. 7c).

**Figure 7.**
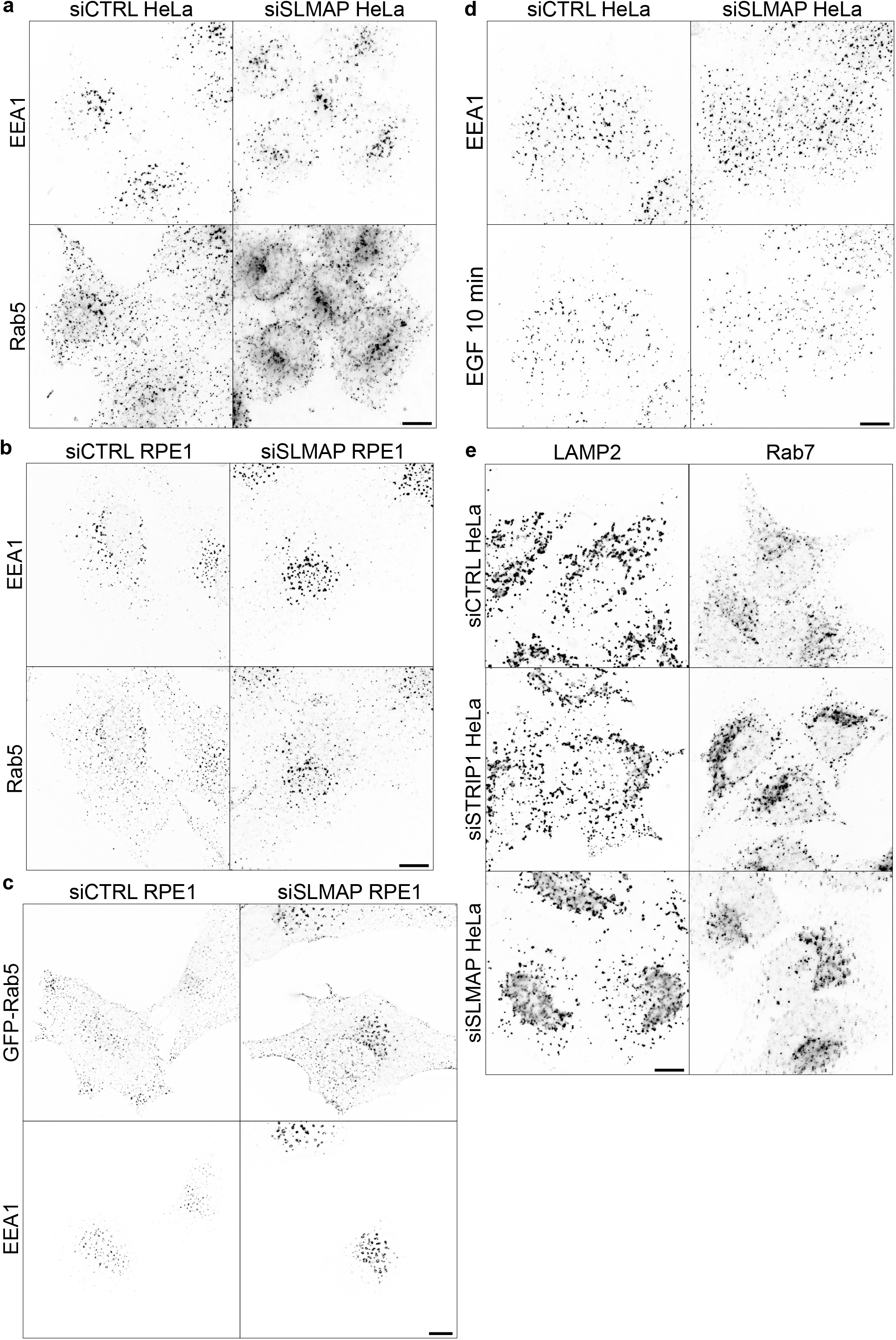
SLMAP is required for endolysosomal organisation. Cells were depleted using siRNAs as indicated and stained for EEA1 and Rab5 (**a,b**), transduced with GFP-Rab5 and stained for GFP and EEA1 (**c**), bound and chased for 10min with ^A88^EGF and stained for EEA1 (**d**), or stained for LAMP2 or Rab7 (**e**). Scale bars = 10 µm.

Overall, these observations on the steady-state behaviour of EEs suggest that SLMAP might have apposing effects to Strip1. We therefore assessed the impact of SLMAP depletion on the formation of new, EGFR-containing EEs. SLMAP depletion did not affect EGFR levels (Fig. S3a). Cells were incubated at 4°C to bind Alexa^488^-EGF, followed by warming cells to initiate EGF uptake. Under these conditions, Alexa^488^EGF had concentrated in EEA1-positive endosomes, many located in the perinuclear region, within 10 min in control cells. However, this concentration of Alexa^488^EGF in EEs was retarded by SLMAP depletion, and EEs were more dispersed than in control-depleted cells (Fig. 7d). Hence, SLMAP depletion appears to retard the formation of new EEs containing internalised EGFR, despite EEA1-containing EEs containing more Rab5 at steady state.

Our data suggest that Strip1 and SLMAP are important for EE behaviour, though generating distinct phenotypes. To explore their impact on later endocytic compartments, we first examined how their depletion affected the distribution of lysosomes. In HeLa cells depleted of Strip1, LAMP2 compartments were larger and more abundant than in control-depleted cells (Fig. 7e). However, the effect of SLMAP depletion on LAMP2 distribution was more striking, with clouds of LAMP2 compartments clustered in the perinuclear region (Fig. 7e). We next explored impacts on Rab7 distribution, since alterations in Rab5 levels caused by depleting STRIPAK may have secondary consequences for Rab7 production. Control-depleted HeLa cells stained strongly for Rab7, with labelling broadly distributed over the cytoplasmic area including on punctate compartments that sometimes concentrated towards the nucleus (Fig. 7e). In Strip1-depleted cells, Rab7 staining on compartments was substantially more intense (Fig. 7e), whereas SLMAP depletion redistributed Rab7 into a dense area close to the nucleus (Fig. 7e), broadly similar to the distribution of LAMP2.

Finally, we explored the impact of SLMAP depletion on lysosomal trafficking in RPE-1 cells. SLMAP depletion strongly reduced the normally rapid uptake of A^448^EGF into larger EEA1-positive endosomes that occurs within 10min (Fig. 8a). Even after 20min, the uptake of Alexa^488^EGF into EEA1-positive endosomes was markedly reduced by SLMAP depletion (Fig. 8a). To follow Alexa^488^EGF trafficking to lysosomes, cells were incubated with lysosomal protease inhibitors during a 60min chase. In control cells, Alexa^488^EGF clearly co-localised with LAMP2-positive lysosomes, whereas Alexa^488^EGF was virtually absent from lysosomes in SLMAP-depleted cells (Fig. 8b). Hence, SLMAP is needed for efficient trafficking of EGFR through the endocytic pathway to lysosomes.

**Figure 8.**
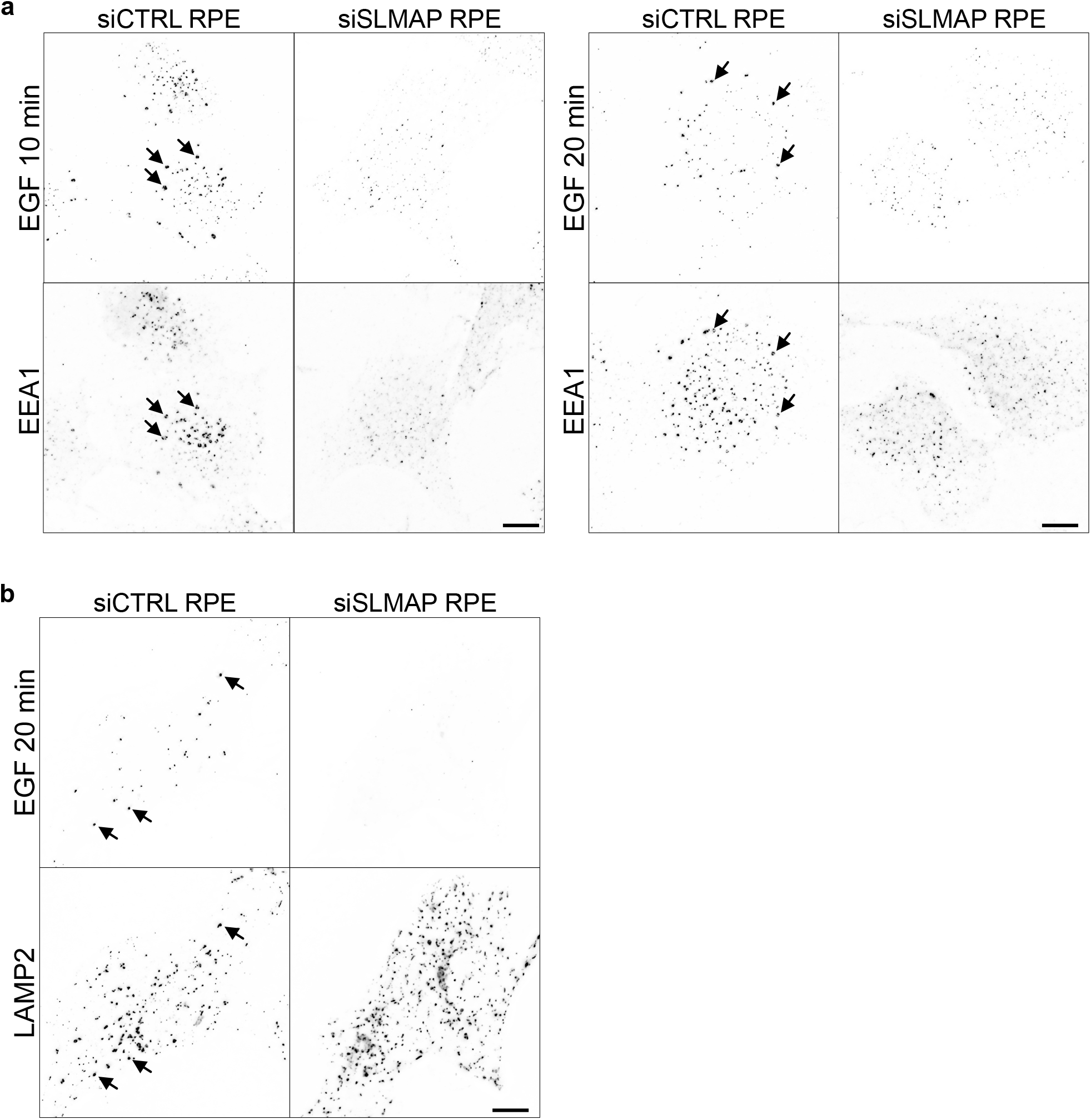
SLMAP is required for lysosomal trafficking of EGFR. **a:** RPE-1 cells were depleted using siRNAs as indicated and pulse-chased with Alexa^488^EGF for the indicated times prior to fixation and labelling for EEA1. **b:** as for a, except cells were incubated with leupeptin and pepstatin to inhibit lysosomal proteases and labelled for LAMP2. Scale bars = 10 µm.

In summary, we have identified a new association between the STRIPAK complex and Rabex5-Rabaptin5 and shown that STRIPAK influences Rab5-dependent endosome biogenesis and EGFR uptake into EEs.

## Discussion

Here we describe the impact of STRIPAK on Rab5-containing EEs. Our starting point was to explore the biochemical impact on the Rab5 GEF Rabex5-Rabaptin5 caused by siRNA-mediated depletion of the ESCRT scaffold HD-PTP. HD-PTP directs MVB formation by interacting with multiple ESCRT complexes, including the ubiquitin-binding ESCRT-0 and ESCRT-I and the membrane-sculpting ESCRT-III subunit CHMP4B (Ali et al., 2013; Doyotte et al., 2008; Stefani et al., 2011). HD-PTP depletion prevents MVB sorting and ubiquitinated MVB cargo instead builds up on the limiting membrane of aberrant EEs (Ali et al., 2013). EEs fail to mature properly to LEs, and ligand-activated EGFR accumulates in clustered, enlarged EEs that contain high levels of Rab5 (Parkinson et al., 2021). Rabex5 binds to and is allosterically activated by ubiquitinated cargo (Lauer et al., 2019; Mattera et al., 2006), and thus the accumulation of this cargo on EEs might partly explain why Rab5 levels remain high when MVB sorting is disrupted (Ott et al., 2025). However, the direct binding of HD-PTP to Rabaptin5 in a manner that is mutually exclusive with HD-PTP binding ESCRT-0 and ESCRT-III (Parkinson et al., 2021) indicates a more direct, mechanistic coupling between MVB sorting and endosome maturation is also at play.

Our MS-based interactomics of Rabaptin5 IPs underscores the hyperactivation of Rab5 in HD-PTP-depleted cells, since we observed increased association of Rab5. This may be a consequence of more Rab5 engaging with the catalytic site within Rabex5 but might also reflect increased Rab5 binding to the Rab5 binding domain at the C-terminus of Rabaptin5. Consistent with the latter possibility, we also observed enhanced association of EEA1 with Rabaptin5 IPs from HD-PTP depleted cells. Rabaptin5 and Rab5 combine in a positive feedback loop to generate endosomal membrane domains rich in Rab5-GTP that are also enriched in EEA1 (Cezanne et al., 2020; Zerial and McBride, 2001). These domains contain high levels of PI3P by virtue of Rab5-GTP recruiting the PI3-kinase VPS34 (Christoforidis et al., 1999b). Although we did not observe enrichment of VPS34 in Rabaptin5 IPs from HD-PTP-depleted cells, there was increased STAM and Hrs, subunits of ESCRT-0 (Raiborg et al., 2002; Ren et al., 2009). ESCRT-0 is recruited to PI3P-rich membrane domains via the FYVE domain of Hrs (Raiborg et al., 2001). In summary, the findings of our unbiased biochemistry are consistent with the accumulation on EEs of Rab5-rich membrane domains in HD-PTP-depleted cells by virtue of hyperactivation of Rabex5-Rabaptin5. We also observed enrichment of the HD-PTP paralogue, Alix/PDCD6IP (Table S1). While we previously found selective binding of HD-PTP but not Alix to Rabaptin5 (Parkinson et al., 2021), these data may point towards a functional interaction between HD-PTP and Alix during EE morphogenesis (Bissig and Gruenberg, 2014; Sun et al., 2015).

By far the most striking alteration in Rabaptin5 interactions in HD-PTP-depleted cells was the increased association of STRIPAK. Nearly all STRIPAK components were identified by MS in Rabaptin5 IPs, including PP2Aase subunits (PP2AA and PP2AC), scaffold proteins (striatins-1, -3, and -4), adaptors (Strip1/FAM40A, CCM3/PDCD10, MOB4, SIKE1, and FGFR1OP2/SIKE2), and the membrane-anchored adaptor, SLMAP (Couzens et al., 2013; Duhart and Raftery, 2020; Hwang and Pallas, 2014; Jeong et al., 2021; Kück and Pöggeler, 2024; Kuck et al., 2019; Tanti et al., 2023; Wigle et al., 1997). Our data suggest that STRIPAK engages Rabex5-Rabaptin5 via more than a single interaction, since exogenously expressed Rabaptin5 engages strongly with SLMAP whereas exogenously expressed Rabex5 engages with Strip1 and SIKE most strongly. While we do not yet know if these interactions are direct, our data suggest that STRIPAK may engage in sequential regulation of Rabex5-Rabaptin5 activity, in keeping with the sequenced assembly of STRIPAK (Niphadkar et al., 2024; Pracheil and Liu, 2013). This would be consistent with the differential effects of Strip1 and SLMAP depletion on EEs that we observe.

Strip1 has previously been linked to Rab5-GEF activity, since *Drosophila Strip* interacts with *Sprint* (Sakuma et al., 2014), a Rab5-GEF related to mammalian Rin1, which acts at an earlier point in the endocytic pathway to Rabex5-Rabaptin5 (Balaji et al., 2012). While the detailed molecular impact of *Strip* on *Sprint* is uncertain, loss of *Strip* phenocopies expression of dominant-negative Rab5 and is rescued by ectoptic Rab5 expression, while *Strip* mutants have reduced levels of *Sprint* protein (Sakuma et al., 2014). Hence, *Strip* is a positive regulator of *Sprint*. Additionally, *Strip* was found to associate with the p150^Glued^ subunit of the dynein regulator dynactin, and endosome localisation was affected in *Strip* mutants (Sakuma et al., 2014). However, this study concluded that these connected functions of *Strip*/Strip1 during EE morphogenesis occur outside of the STRIPAK complex, in contrast to the conclusions of our study. Strip1 associates with the catalytic Vps9 domain (Sakuma et al., 2014) of *Sprint*, which is conserved across Rab5-GEFs including Rabex5 (Carney et al., 2006; Delprato and Lambright, 2007), and hence binding to the Rabex5 Vps9 domain may underlie the association between Strip1 and Rabex5. Depletion of Strip1 reduces cellular levels of Rab5-GTP without affecting Rab5 protein levels, and broadly phenocopies Rabaptin5 depletion. However, we do not yet know if Strip1 directly affects Rabex5 activity, and whether this effect might occur through allosteric mechanisms (Lauer et al., 2019) or by altering the membrane recruitment of Rabex5-Rabaptin5 (Kalin et al., 2015). While Strip1 depletion closely phenocopies Rabaptin5 depletion, other Vps9 proteins likely contribute to some of the effects on EEs that we observe.

STRIPAK PP2Aase activity has many cellular roles, notably including the deactivation of protein kinases(Kück and Pöggeler, 2024). Hippo kinase an exemplar, with Hippo activated by phosphorylation within its linker region. SLMAP binds to phospho-peptides within this region, thereby recruiting Hippo to STRIPAK PP2Aase and leading to Hippo inactivation (Bae et al., 2017; Zheng et al., 2017). Other MST kinases follow this pathway, including MST4/STK26, and we find STK26 among HD-PTP depletion-enhanced Rabaptin5 interactors. Given that Rabaptin5 is subject to phosphorylation to control its activity (Christoforides et al., 2012), and that Rabaptin5 is hyper-phosphorylated in HD-PTP-depleted cells (Parkinson et al., 2021), it is possible that the recognition of Rabaptin5 by STRIPAK is connected to Rabaptin5’s phosphorylation status.

Through its activity towards Hippo and other protein kinases, STRIPAK has been linked to coordination of cell growth signalling and stress response pathways (Ardestani and Maedler, 2020; Bae et al., 2020; Honda et al., 2023; Kim et al., 2020; Li et al., 2023; Seo et al., 2020; Zhong et al., 2024). However, STRIPAK also has roles in polarised membrane trafficking (Kean et al., 2011; Lant et al., 2015) and emerging roles in autophagy (Kück and Pöggeler, 2024). STRIPAK controls the movement of autophagosomes in *Drosophila* neurons, via interaction of striatin (CKA) with ATG8 and dynein. STRIPAK also regulates autophagy in muscle cells by associating with Stv and NAUK kinases (Guo et al., 2023), while STRIPAK acts in conjunction with MAP4K4 to regulate autophagy in accordance with cellular stress (Seo et al., 2024). Furthermore, crosstalk between ULK1 and STRIPAK (Hu et al., 2021; Siva Sankar et al., 2022) highlights the interplay between STRIPAK signalling and the initiation of autophagosome formation. Our work now helps place STRIPAK at the centre of early endosome biology.

## Materials and methods

### Reagents

#### Antibodies

The following commercial antibodies were used: Mouse: anti-HA (GenScript; Cat. A01244; batch A204810; 1:2500 for WB); anti-FLAG (Proteintech; Cat. 66008-4-Ig; Lot 10027647; 1:1000 for IF); anti-GFP (Roche; Cat. 11814460001; Lot 12600500 1:1000 for IF); anti-EEA1 (BD Biosciences; 14/EEA1; Cat. 610457; batch 6302827; 1:400 for IF); anti-LAMP2 (Developmental Studies Hybridoma Bank; Cat. H4A6; 1:400 for IF); anti-Rab5A (Proteintech; Cat. No. 66339-1-Ig; Lot No. 10003870; 1:400 for IF); anti-Rab5 (BD Biosciences; Cat. 610725; batch 11971; 1:2500 for WB); anti-SLMAP (AbNova; Cat. H00007871-M08; Lot. No. N5291-2A7; 1: 1000 for WB); anti-Striatin (BD Biosciences; Cat. 610838; Lot. No. 2137043; 1:1000 for WB); anti-His_6_ clone H1029 (Sigma; Cat. 062M4809; 1:5000 for WB); anti-vinculin (hVIN1 Merck Cat. No. V9131 Lot. No. 0000128055; 1:5000 for WB). Rabbit: anti-HD-PTP (Proteintech; Cat. 10472-1-AP; lot 00018154; 1:5000 for WB); anti-EEA1 (Cell Signalling Technologies; C45B10; Cat. 3288S; batch 8; 1:400 for IF); anti-Rabex-5 (Thermo Scientific - Invitrogen; Cat. PA5-21117; batch RF2221849; 1:2500 for WB) anti-Rabex-5 (Proteintech; Cat. No. 12735-1-AP; Lot. No. 00021439; 1:1000 for WB); anti-FAM40A (AbCam; Cat. No. AB199851; Lot. No. 1021914; 1:2500 for WB); anti-Rabaptin-5/RABEP1 (Proteintech; Cat. 14350-1-AP; batch 00005401; 1:5000 for WB, 1:200 for IF); anti-Rab5 clone C8B1 (Cell Signalling Technologies; Cat. 3547S; batch 7; 1:500 for IF; 1:2500 for WB); anti-PPP2R1 (Abcam; Cat. No. ab24736; Lot No. GR3321194-1; 1:1000 for WB); anti-Rab7 (Proteintech; Cat. No. 84741-1-RR; Lot. No. 23012663; 1:400 for IF); anti-SLMAP (Proteintech; Cat. No. 25220-1-AP; Lot. No. 00022209; 1:000 for WB); anti-GAPVD1 (Abcam; Cat. ab86033; batch GR265707-8; 1:5000 for WB);. Goat: anti-GST (GE Healthcare - Cytiva; Cat. 27457701V; batch 9528344; 1:10000 for WB); anti-HA (GenScript; Cat. A00168; batch A102608; 1:2500 for WB; 1:100 for IF). Fluorescent secondary antibodies for microscopy and Western blotting were from Jackson ImmunoResearch Laboratories (PA, USA): Fluorescent secondary antibodies were from Jackson ImmunoResearch Laboratories (PA, USA): (Alexa Fluor 680-conjugated AffiniPure Donkey Anti-Rabbit IgG; Cat. 711-625-152; batch 139907; 1:5000 for WB); (Alexa Fluor 680-conjugated AffiniPure Donkey Anti-Mouse IgG; Cat. 715-625-150; batch 145146/153931; 1:5000 for WB); (Alexa Fluor 790-conjugated AffiniPure Donkey Anti-Rabbit IgG; Cat. 711-655-152; batch 132235; 1:5000 for WB); (Alexa Fluor 790-conjugated AffiniPure Donkey Anti-Goat IgG; Cat. 705-655-147; batch 143129; 1:5000 for WB); (Alexa Fluor 680-conjugated AffiniPure Donkey Anti-Rabbit IgG; Cat. 711-625-152; batch 149402; 1:5000 for WB); (Alexa Fluor 680-conjugated AffiniPure Donkey Anti-Goat IgG; Cat. 705-625-147; batch 133233; 1:5000 for WB); (Alexa Fluor 488-conjugated AffiniPure Donkey Anti-Mouse IgG; Cat. 715-545-151; 1:800 for IF); (Alexa Fluor 594-conjugated AffiniPure Donkey Anti-Mouse IgG; Cat. 715-585-151; 1:800 for IF); (Alexa Fluor 488-conjugated AffiniPure Donkey Anti-Rabbit IgG; Cat. 711-545-152; 1:800 for IF); (Alexa Fluor 594-conjugated AffiniPure Donkey Anti-Rabbit IgG; Cat. 711-585-152; 1:800 for IF).

#### Oligonucleotides

All siRNA oligonucleotides are listed in Table S3. Allstars negative control siRNA (Qiagen) was used for non-silencing conditions.

### DNA

#### Mammalian expression vectors

GFP-Rabaptin5 was used as reported (Parkinson et al., 2021). FLAG-STRIPAK and HA-Rabex5 plasmids were generated using templates purchased from Sino Biological using standard PCR based cloning techniques.

### Cell culture, transfections and treatments

HeLa (ATCC CCL-2) and HEK293T (ATCC CRL-3216) cells were grown in DMEM supplemented with 1% NEAA, 10% Foetal Calf Serum (HyClone; Cytiva) and 2mM glutamine (Sigma) at 37°C and 8% CO_2_. hTERT-RPE-1 cells were grown in DMEM/F-12 supplemented with 1% NEAA, 10% Foetal Calf Serum (HyClone; Perbio) and 2mM glutamine (Sigma) at 37°C and 5% CO_2_. Cells were verified and checked routinely for contamination. HeLa and hTERT RPE-1 DNA transfections were performed using JetPEI (Polyplus) or Lipofectamine 2000 (Invitrogen). HEK293T DNA transfections were performed using Branched PEI Mw - 25kDa (Sigma): briefly, complexes of DNA:PEI (1 mg/ml) at a ratio of 1:3 were formed and cells were analysed 20-24 h post-transfection. Transfection with siRNA duplexes (Table S3) was performed using INTERFERin (Polyplus) according to the manufacturer’s instructions. Cells were analysed 16-24 h post-transfection. GFP-Rab5 lentivirus was a gift from Martin Lowe (University of Manchester). Supernatant containing lentiviral particles was used to transduce cells for 3 days prior to fixation

### Generation of Strip1-mAID-FKBP12 double degron cell lines

To generate Strip1-mAID-FKBP12 double degron (Strip1-DD) cell lines, the mAID-FKBP12 double degron was inserted at the C-terminus of Strip1 using two guide sequences (GCAGTGAGGCTGTTGGTTAG and TGTTGGTTAGGGGACTGAAA) targeting the C terminus of Strip1. Guide sequences were introduced into RPE-1 cells in which OsTIR1(F74G) and the zeocin resistance gene were simultaneously inserted into the safe harbour AAVS site, using a plasmid encoding a modified Cas9 in which the C-terminus of the eSpCas9 enzyme is fused to amino acids 1–110 of human Geminin. This plasmid (esPcas9-hGeminin-AAVS-Strip1) contained a guide to the AAVS safe harbour site as well as guides directed to Strip1 (Belbelazi et al., 2026). Screening of clonal lines was performed by PCR using a Bioline Isolate II genomic DNA kit, and KOD polymerase (Merck) in accordance with manufacturer’s instructions and the primers: Fwd: 5’-CTCTGTGTGTGCGTGTTGCTC-3’, Rev: 5’-GCCTCTGAAAGCCAAAAGGAAAA-3’. Sanger sequencing was performed to confirm the presence of the tags (Genewiz). Strip1-DD was depleted using 1 uM 5-PH-IAA (MedChemExpress, HY-134653) or 0.1 uM dTAGv1 (MedChemExpress, HY-145514D) for 4h. dTAGV-1 was used as routine.

### SILAC labelling

For quantitative mass spectrometry, HeLa cells were labelled in SILAC RPMI (PAA Laboratories GmbH, Germany) supplemented with 10% dialyzed fetal bovine serum (Sigma), 2 mM glutamine (Gibco), 100 U/ml penicillin and 100 μg/ml streptomycin for 15 days to ensure complete incorporation of amino acids, which was verified by MS analysis. Two cell populations were obtained: samples labelled with natural variants of the amino acids (light label; Lys0, Arg0), which were used as control samples, and samples labelled with heavy variants of the amino acids (heavy label; L-[13C6,15N4]Arg (+10) and L-[13C6,15N2]Lys (+8); Lys8/Arg10) which we used for cells depleted of HD-PTP. The light amino acids were from Sigma and the heavy variants were from Cambridge Isotope Labs (Massachusetts, US).

### Sample preparation for total proteome and IP

SILAC Light/Heavy cells (Light cells were transfected with Control siRNA, Heavy cells were transfected with HD-PTP siRNA; an additional batch of Light cells were transfected with Control siRNA and used as control for the IPs; 8 x 15cm dishes per condition) were transfected with siRNA for 48h. Cells were washed 3x in PBS and lysed in IP buffer (25mM Tris-HCl pH7.5, 150mM NaCl, 0.5m MEDTA, 0.5% IGEPAL, 2 mM NaVO_4_, 50mM β-glycerophosphate, 50mM NaF, 10mM N-ethylmaleimide, 1mM PMSF, 1x EDTA-free protease inhibitor cocktail (Roche); 1ml per dish), agitated for 30min at 4°C and centrifuged for 30min at 14,000rpm in a benchtop microfuge. A fraction (20 µl) of the input from each dish was combined for total proteome measurement and Light and Heavy samples mixed 1:1. The remainder of the input was combined and used to perform native Rabaptin5 immunoprecipitation, and Light and Heavy samples mixed 1:1 after IP washes. Immunoprecipitation was performed using rabbit anti-RABEP1 antibody (Proteintech). Briefly, combined samples for each condition were incubated with rotation with rabbit anti-RABEP1 and M-280 sheep anti-rabbit Dynabeads (ThermoFisher) overnight at 4°C. Beads were washed 5x in lP buffer, then once in PBS before eluting in sample buffer. A fraction of each eluant was removed for validation and the remainder combined for proteomics.

### Sample preparation for quantitative proteomics

Total proteome and native Rabaptin5 IP samples were processed similarly using in-gel digestion. Samples were prepared in sample buffer containing 10 mM DTT, alkylated with 5.5 mM CAA and run on 1.00 mm Invitrogen NuPAGE 4–12% Bis-Tris Gel (Invitrogen) in NuPAGE MOPs running buffer (Invitrogen), as previously described (Francavilla et al., 2016). Briefly, gels were washed in ddH2O, and proteins were subsequently fixed and stained with Colloidal Coomassie Stain (Invitrogen). Each sample was equally separated into four fractions, and de-stained using 50% 20 mM ammonium bicarbonate (NH_4_HCO_3_) + 50% ethanol (EtOH). Peptides were digested by trypsin in 50 mM NH_4_HCO_3_, neutralized and extracted using increasing concentrations of acetonitrile (ACN), starting with 50% CAN, 3% trifluoroacetic acid (TFA) and ending with 100% ACN. Digested peptides were evaporated to a final volume of 100 μL in a speed vacuum and loaded on C18 STAGE-tips (Francavilla 2016). Peptides were eluted from STAGE-tips with 40% ACN followed by 60% ACN.

### Mass spectrometry analysis

Purified peptides were analysed by LC-MS/MS using an UltiMate® 3000 Rapid Separation LC (RSLC, Dionex Corporation, Sunnyvale, CA) coupled to a Orbitrap Exploris 480 (Thermo Fisher Scientific, Waltham, MA) mass spectrometer (Watson et al., 2022). Mobile phase A was 0.1% formic acid (FA) in water and mobile phase B was 0.1% FA in ACN and the column was a 75 mm × 250 μm inner diameter, 1.7 mM CSH C18, analytical column (Waters). Samples were taken up and transferred to the 5 μl loop and loaded onto the column at a flow of 500 nl/min at 1% B for 4 min. The loop was then taken out of line and the flow was reduced from 300 nl/min in 1 min. Peptides were separated using a gradient that went from 1 % to 6 % B in 3 min, from 6% to 18% B in 67 min, from 18% B to 29% B in 11 min, and finally from 29% to 65% in 1 min for the column was which was sustained for 6 min, ending the run with an equilibration step by going down to 2% B in 1 min and sustaining it for the rest of the run at 100 min. Mass spectrometry data were acquired in a data-dependent (DDA) manner for 100 min in positive mode. Peptides were selected for fragmentation automatically by data-dependent analysis on the basis of the top 12 with m/z between 300 to 1750 Da at a minimum intensity of 5000 and a charge state of 2, 3 or 4 with a dynamic exclusion set at 15 s and a normalized AGC target of 300%. The MS Resolution was set at 120,000 with an AGC target of 3e6 and a maximum fill time set at 20 ms. The MS2 Resolution was set to 30,000, with a normalised AGC target, and a maximum fill time of 45 ms. The isolation window was 2.6 Da and the relative collision energy for HCD fragmentation was 28%.

### FragPipe search

Protein identification and quantification were performed in FragPipe (v24.0) using the built-in SILAC3 workflow, using two-channel (light/heavy) SILAC as described below. Spectra were searched with MSFragger against the UniProt *Homo sapiens* proteome (taxonomy ID 9606; downloaded 04/09/2026), with common contaminants and reverse decoys. Strict trypsin specificity (cleavage C-terminal to K/R) was used, allowing up to two missed cleavages and peptide lengths of 7–40. Precursor and fragment mass tolerances were ±20 ppm, with mass calibration and parameter optimization enabled and isotope errors of 0/1/2 considered. Carbamidomethylation of cysteine (+57.02146 Da) was set as a fixed modification. Variable modifications were methionine oxidation (+15.9949 Da), protein N-terminal acetylation (+42.0106 Da), and heavy lysine (+8.014199 Da) and arginine (+10.008269 Da), with a maximum of five variable modifications per peptide. PSMs were rescored with MSBooster using predicted retention times and fragment spectra (DIA-NN models) and Percolator, followed by protein inference with ProteinProphet. Results were filtered to 1% FDR at the PSM, peptide and protein levels using Philosopher. SILAC quantification was performed with IonQuant using light (K0, R0) and heavy (K8.014199, R10.008269) channels, an m/z tolerance of 10 ppm, a retention time tolerance of 0.4 min, a minimum of two isotopic peaks and three scans per feature, with re-quantification and normalization enabled. Match-between-runs was enabled (RT tolerance 1 min, ion FDR 1%, top 10 runs).

### MS Data Analysis

Downstream data analysis was performed using R in RStudio and a Quarto Markdown document using the following packages: renv, tidyverse, Msnbase, limma, clusterProfiler, org.Hs.eg.db, forcats, ggrepel, UpSetR, Ggally. All software and package versions can be found in the GitHub repository referenced in the Data Availability section.

For the IP dataset, data was filtered for contaminants and missing values based on the *Median Log2 Ratios HL* values, Different normalization techniques were compared (median normalization, quantile normalization, normalization to the bait), and normalization to the bait was chosen based on the quantitative changes of known interactors. To assess differentially abundant interactors (per unit of bait) limma was used using eBayes, then Gene Ontology (GO) Enrichment was performed on the significant hits. All figures related to the IP dataset in the Manuscript were made using ggplot2, except for the sample-sample correlations, where Ggally was used.

For the Proteome dataset, filtering was performed similarly to the IP dataset, then QC steps were performed to judge general data quality such as quantification depth, overlap between identifications, and similarity between replicates. Then, proteins found *significant* in the IP dataset were plotted along with the bait protein, to assess whether the differences in IP could be attributed to a difference in protein expression. Data was quantile normalized, and differentially abundant proteins were identified using limma and eBayes, and GO enrichment was performed on significant hits. All figures related to the Proteome dataset in the Manuscript were made using ggplot2, except for the sample-sample correlations, where Ggally was used.

### Immunofluorescence, imaging and analysis

Cells were fixed in 3% formaldehyde for 10 min and quenched with 0.1M glycine, 50 mM Tris-HCl pH 7.5 for 10 min, then permeabilised for 3-5 min in PBS containing 0.1% Triton X-100. For some antibody combinations, cells were fixed in methanol at -20°C. To follow EGF uptake, cells were incubated for 3h at 37°C in serum-free DMEM at 8% CO_2_, then incubated for 60min on ice with 100ng/ml Alexa^488^streptavidin conjugated to EGF-biotin (Alexa^488^EGF; Thermo Fisher) in binding medium (CO_2_-independent medium containing 2mg/ml BSA). Cells were then chased at 37°C in serum-free DMEM at 8% CO_2_ before fixing in formaldehyde and processing for IF. To follow Alexa^488^EGF transport to lysosomes, cells were chased in the presence of 50µM leupeptin, 50 µM pepstatin to inhibit lysosomal degradation.

Standard fluorescence experiments were imaged on an Olympus BX51 upright microscope fitted with a 60 x 1.4 NA Plan Apo objective and CoolSnap ES camera, and 12-bit images captured using MetaVue software. Confocal microscopy was carried out using a Leica SP8 Lightening confocal microscope. Alternatively, images were acquired using an Olympus IX83 inverted microscope using Lumencor LED excitation, and a 60x/ 1.42NA Plan Apo objective. Images were collected using a R6 (Qimaging)] CCD camera with a Z optical spacing of 0.2μm. Raw images were then deconvolved using the Huygens Pro software (SVI) and maximum intensity projections of deconvolved images were generated using ImageJ. All images were processed and analysed using ImageJ. All comparison images were acquired using the same acquisition settings and adjusted using near-identical brightness/contrast settings. Antibody staining intensity over endosomes was measured on wide-field captured images using ImageJ. For HD-PTP KDs, to quantify the percentage of cells with normal or aberrant endosome distributions, conditions were blinded and the proportion of normal cells in randomly selected fields was assessed by eye: 100 cells were scored for each condition; any cells that did not exhibit enlarged or clustered endosomal morphology were considered normal.

### Protein extraction and Western blotting

To perform Western blotting of total cell extracts, cells in dishes were washed 3 times in PBS, and then lysed in 2x SDS PAGE sample buffer. Samples were run on polyacrylamide gels and transferred to PVDF membranes (Millipore). Blots were scanned with a Li-Cor Bioscience Odyssey scanner. Quantitation was performed using ImageStudio software.

### Coimmunoprecipitations and pull-downs

For native IPs, cells were lysed in IP buffer (25mM Tris-HCl pH7.5, 150mM NaCl, 0.5% (v/v) IGEPAL CA-630, 1x phosphatase inhibitor cocktail 2 and 3 (Sigma), 1x Protease Inhibitor cocktail EDTA-free tablet (Roche)). Lysates were clarified at 14,000 rpm for 30 min at 4°C and incubated with antibodies at 4°C for at least 4 h, then 1-2 h with 10µl protein A or protein G Sepharose slurry (Zymed), pre-blocked with 50 mg/ml BSA or 10ul Flag Agarose (Sigma). Beads were washed three times in IP buffer, then analysed by Western blotting (WB). Panels for all figures have been cut from the same exposures.

### GST–APPL1N and GST pull-down assay

GST–APPL1N and GST control proteins were expressed in Rosetta (DE3) cells transformed with pGEX–APPL1N or pGEX-4T-3, respectively. Protein expression was induced with 0.4 mM IPTG at 18 °C overnight. Bacterial pellets were resuspended in PBS-plus (PBS containing 1% Triton X-100, 1 mM DTT, 1 mM EDTA, and protease inhibitor cocktail tablets as above), lysed by sonication, and clarified by centrifugation at approximately 20,000 × g for 30 min at 4°C. Supernatants containing GST proteins were stored at −80 °C until use. Glutathione Sepharose 4B beads (15 µL slurry per sample) were washed with PBS-plus and incubated with 300 µL bacterial lysate containing GST or GST–APPL1N for 60 min at RT. Protein-loaded beads were washed five times with lysis buffer (25 mM HEPES-KOH, pH 7.4, 100 mM NaCl, 5 mM MgCl_2_ 0.1% Triton X-100, 10% glycerol, and 1 mM DTT). For GST pull-down assays, hTERT-RPE-1 cells were grown in 6-well plates (three wells per sample) and transfected with 20 nM target siRNA using INTERFERin (Polyplus), followed by transfection with 200 ng EGFP–Rab5A and 5.4 µg pBS carrier DNA using jetPEI (Polyplus). Cells were lysed in lysis buffer supplemented with 1x phosphatase inhibitor cocktail 2 and 3 (Sigma), 1x Protease Inhibitor cocktail EDTA-free tablet (Roche), and clarified lysates were incubated with GST- or GST–APPL1N-loaded beads for 1h at 4 °C. Beads were washed five times with lysis buffer, and bound proteins were analysed by SDS–PAGE and Western blotting.

### Experimental design and statistical analysis

All data are presented as mean ± SD unless otherwise stated. Data were first tested for normality using the D’Agonstino-Pearson omnibus K2 normality test to determine the appropriate statistical test. Since all data with large sample size were found to fit a normal distribution, parametric statistics were employed. For experiments with n numbers lower than 8, data was assumed to fit a normal distribution without formal testing. Unpaired two-tailed Student’s t tests were used for statistical comparisons between 2 groups, and for experiments containing 3 or more groups, one-way analysis of variance (ANOVA) was employed. Two-way ANOVAs were used to analyse data affected by two categorical independent variables. To compare all variables with one another, a Bonferroni’s post-hoc test for multiple comparisons was used. To compare each variable to a single control variable, a Dunnett’s post hoc test was used. All statistical tests were performed on GraphPad Prism (v6.0) and differences were considered significant if p˂0.05; *p˂0.05, **p˂0.01, ***p˂0.001.

## Supporting information

Wunderley Supplemental information

## Figure Legends

**Figure S1. QC control of Rabaptin5 IPs for SILAC MS-based interactomics a**: Replicate Rabaptin5 IPs from control- or HD-PTP-depleted cells blotted for Rabaptin5, HD-PTP, or GAPDH as a loading control. A control IP from control-depleted cells using an unrelated IgG is shown alongside replicate 3 to confirm the specificity of the Rabaptin5 IP. **b:** Density curves of HL median SILAC ratios for the 3 biological replicates. **c:** Pairwise sample-sample correlations including density curves in the diagonal and sample-sample scatterplots and Pearson correlation coefficients for the bait-normalised (STRIPAK1) IP samples. **d:** Quality control plot that show the necessity of bait-normalisation highlighting known interactors of RABEP1 according to the IntAct database. Real, known interactors such as RABGEF1 follow the bait (close to 0 difference between IP2 and the mean of the other two replicates), while the background does not (artifacts from high-throughput screenings in the database). **e:** data completeness barplot of the total proteome samples. **f:** Density curves of the total proteome samples showing good overlap. **g:** Pairwise sample-sample correlations including density curves in the diagonal and sample-sample scatterplots and Pearson correlation coefficients for the total proteome samples. **h:** Lineplots of the median HL SILAC ratios of the significant interactors in the total proteome showing close to 0 values indicating largely unchanged protein levels in HD-PTP depleted cells. **i:** Lineplots of the median HL SILAC ratios of the bait protein and the detected components of the STRIPAK complex.

**Figure S2. Rabaptin5β (Rabep2) does not associate with either HD-PTP or STRIPAK a:** Lysates from HEK293 cells co-expressing GFP-Rabaptin5α (Rabep1) or GFP-Rabaptin5β (Rabep2) and HA-HD-PTP(Bro1-CC) were subjected to anti-HA IP. Left: representative experiment. Right quantitation from 3 independent experiments. **b:** GFP-Rabaptin5α or GFP-Rabaptin5β were expressed in HeLa cells with or without HA-HD-PTP(Bro1-CC) and stained with anti-HA or imaged for GFP fluorescence. Bar = 20 µm. **c:** Lysates from HEK293 cells co-expressing GFP-Rabaptin5α (Rabep1) or GFP-Rabaptin5β (Rabep2) and FLAG-SLMAP were pulled-down using GFP-Trap beads and blotted for GFP or FLAG. Left: representative experiment. Right quantitation from 3 independent experiments. Scale bars = 20 µm.

**Figure S3. Impact of STRIPAK depletion on early endosomes. a,b:** Western blots of siRNA depletion experiments. **c:** RPE-1 cells were depleted using siRNAs as indicated and stained for EEA1 and Rab5. **d:** RPE-1 cells were depleted using siRNAs as indicated and transduced with GFP-Rab5 and stained for GFP and EEA1. Scale bars = 10 µm.

## Acknowledgements

We thank the University of Manchester Faculty of Biology Medicine and Health Bioimaging and Bio-MS facilities. This work was supported by the Medical Research Council (G0701140) and the Wellcome Trust (212246/Z/18/Z). CF has received funding from the Wellcome Trust (107636/ Z/15/Z and 107636/Z/15/A) and the Novo Nordisk Foundation Young Investigator Award (call 2022, NNF22OC0070845).

## Author contributions

LW, GP, HS and PW designed and performed biochemistry and imaging experiments; GP and PF performed proteomics experiments; SS analysed proteomics datasets; EWT generated reaents; CF and PGW designed and supervised the project; GP, LW, SS, CF and PGW wrote the paper.

