## Supplementary material for "STRIPAK associates with Rabex5-Rabaptin5 to promote Rab5-dependent early endosome formation and EGFR transport": Wunderley Supplemental information

**Wunderley et al**
**Supplemental information**

**Table S3. siRNA oligonucleotides**

| **Target** | **Sequence** |
|---|---|
| HDPTP (PTPN23) | GCAAACAGCGGAUGAGCAAUU |
| RABEP1 | AAGUAGUAUGCUGUAUGAAUA |
| RABEP2 | GGACAGAGAUGGAGCGGGU<br>GAGGUGAGCGAGAGCACGA<br>ACCAUGAGGUAAAGCGGUU<br>UCGUACUGCCCAUGGAAAA |
| STRIP1 (FAM40A) | CAGGGTCCTGCTCAACATCAT |
| SLMAP | GCAUCUAACCAAAGCGGUA |
| STRN | AAGUGAAGAAUAGUUGAUCGA |
| STRN3 | AAGCAGACUUGACGGUAACAA |
| STRN4 | AAUAUAGGUUCCACAUCUAGC |

### Figure S1

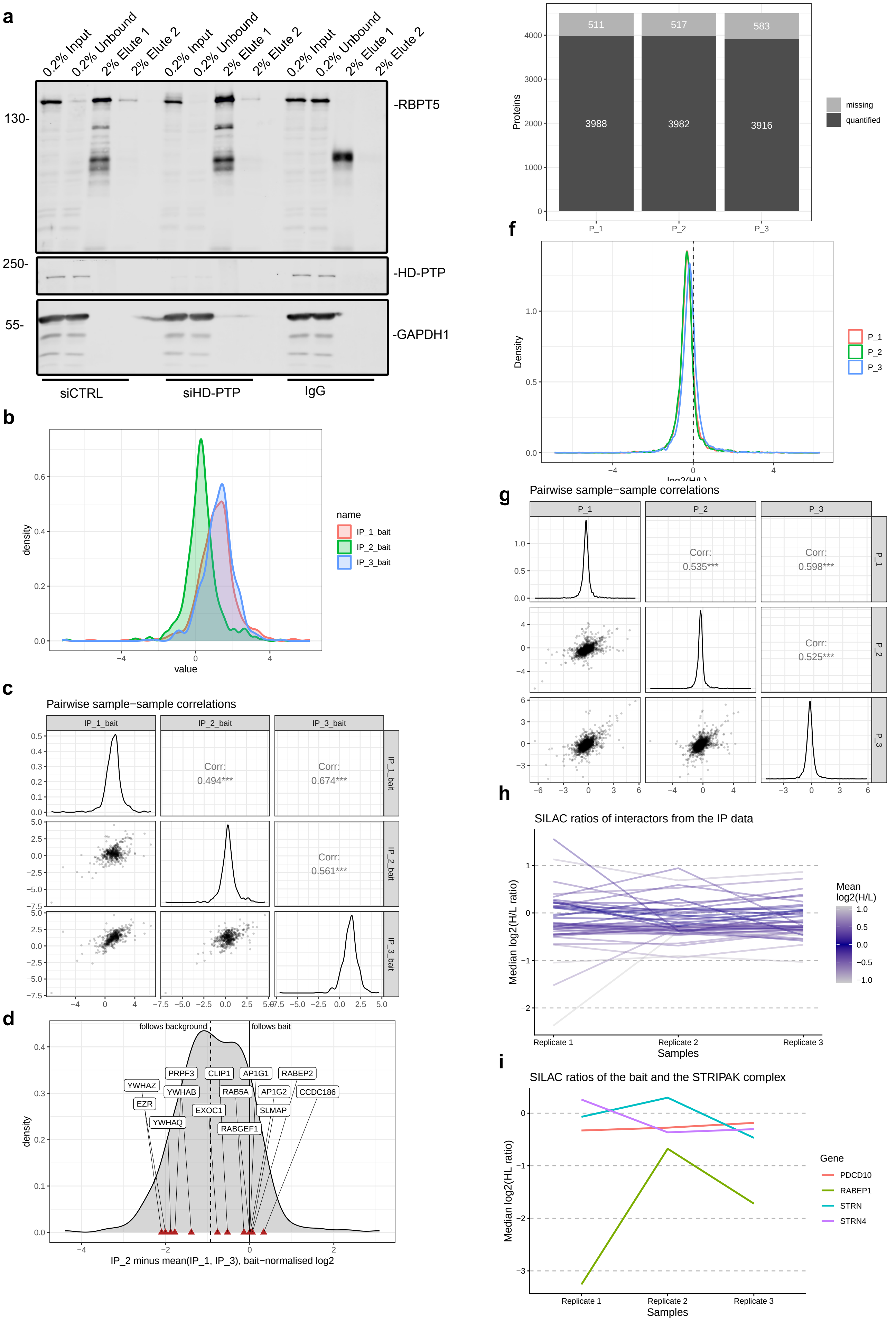

a
0.2% Input
0.2% Unbound
2% Elute 1
2% Elute 2
130-
250-
55-
-RBPT5
-HD-PTP
-GAPDH1
siCTRL
siHD-PTP
IgG
b
density
value
name
IP_1_bait
IP_2_bait
IP_3_bait
c
Pairwise sample−sample correlations
IP_1_bait
IP_2_bait
IP_3_bait
Corr:
0.494***
Corr:
0.674***
Corr:
0.561***
d
follows background
follows bait
YWHAZ
EZR
YWHAQ
PRPF3
YWHAB
EXOC1
CLIP1
RAB5A
RABGEF1
AP1G1
AP1G2
SLMAP
RABEP2
CCDC186
density
IP_2 minus mean(IP_1, IP_3), bait−normalised log2
e
Missing median SILAC ratio values per replicate
Proteins
511
517
583
3988
3982
3916
P_1
P_2
P_3
missing
quantified
f
Density
log2(H/L)
P_1
P_2
P_3
g
Pairwise sample−sample correlations
P_1
P_2
P_3
Corr:
0.535***
Corr:
0.598***
Corr:
0.525***
h
SILAC ratios of interactors from the IP data
Median log2(H/L ratio)
Replicate 1
Replicate 2
Replicate 3
Samples
Mean
log2(H/L)
i
SILAC ratios of the bait and the STRIPAK complex
Median log2(HL ratio)
Replicate 1
Replicate 2
Replicate 3
Samples
Gene
PDCD10
RABEP1
STRN
STRN4

#### Figure S3

a

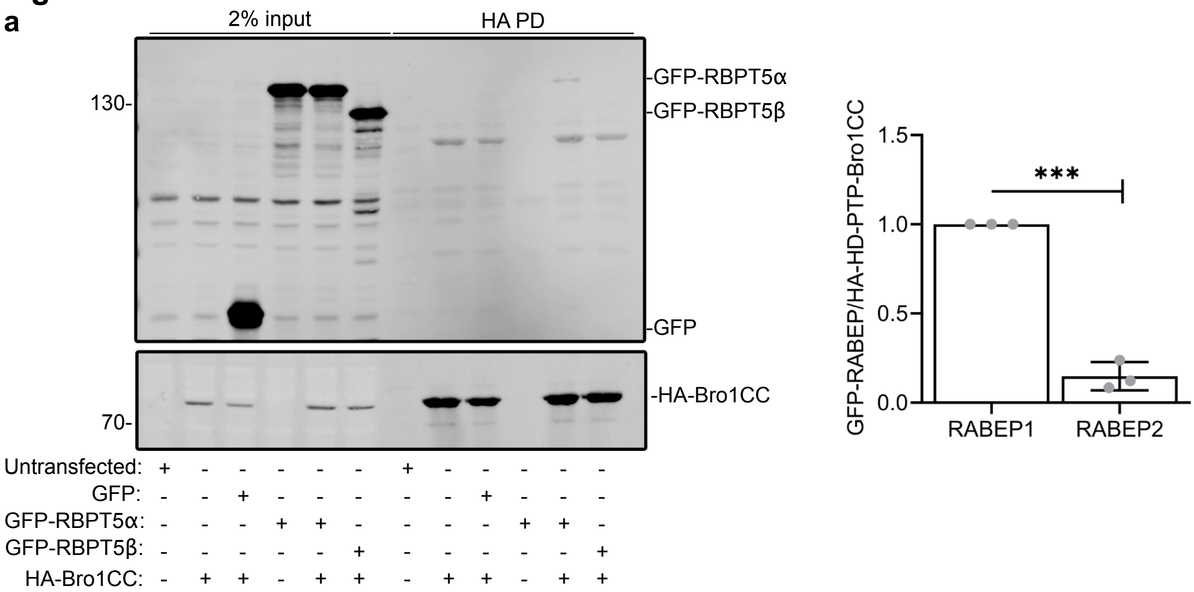


b

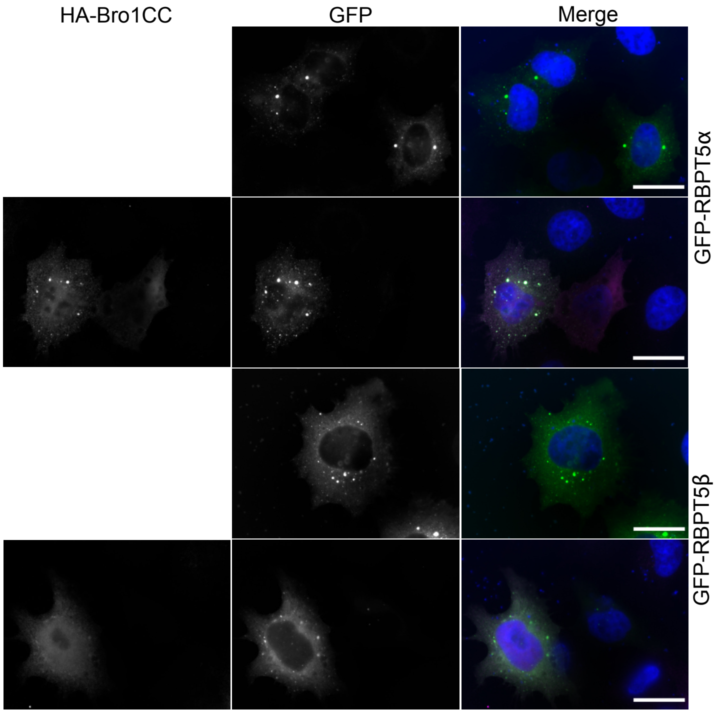


c

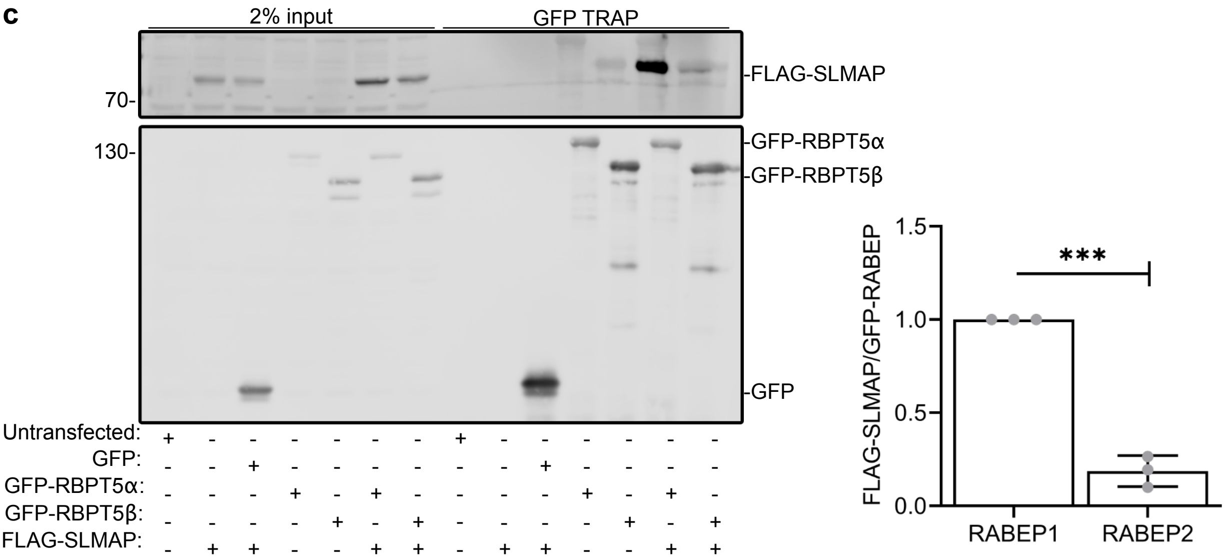

#### Figure S3

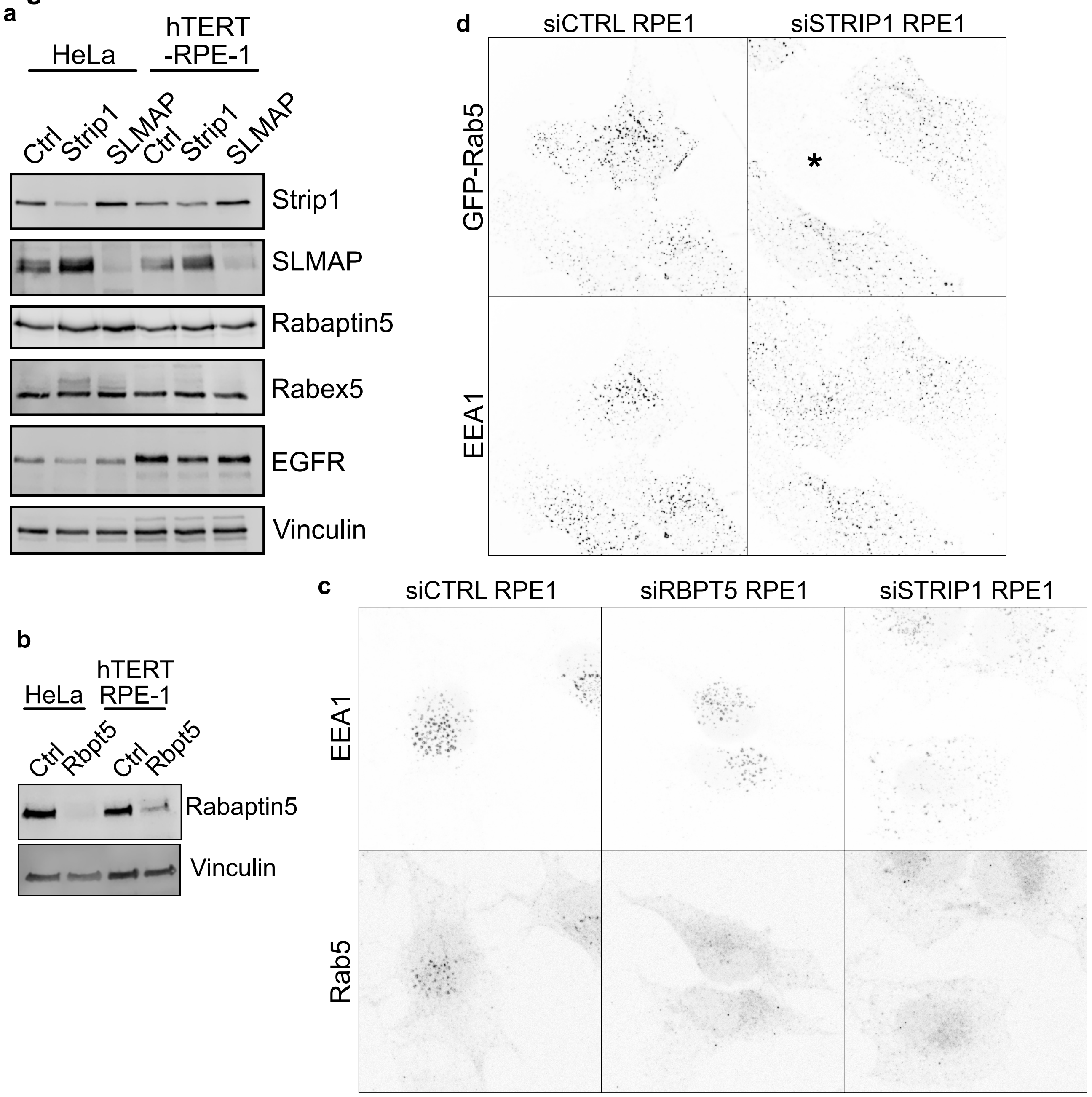
a
HeLa
hTERT -RPE-1
Ctrl
Strip1
SLMAP
Ctrl
Strip1
SLMAP
Strip1
SLMAP
Rabaptin5
Rabex5
EGFR
Vinculin
b
HeLa
hTERT RPE-1
Ctrl
Rbpt5
Ctrl
Rbpt5
Rabaptin5
Vinculin
c
siCTRL RPE1
siRBPT5 RPE1
siSTRIP1 RPE1
EEA1
Rab5
d
siCTRL RPE1
siSTRIP1 RPE1
GFP-Rab5
EEA1
*
